# Nasally-delivered interferon-λ protects mice against upper and lower respiratory tract infection of SARS-CoV-2 variants including Omicron

**DOI:** 10.1101/2022.01.21.477296

**Authors:** Zhenlu Chong, Courtney E. Karl, Peter J. Halfmann, Yoshihiro Kawaoka, Emma S. Winkler, Jinsheng Yu, Michael S. Diamond

**Author notes:** Corresponding author: Michael S. Diamond, M.D., Ph.D. **Lead Contact**: Michael S. Diamond, M.D., Ph.D.

## Abstract

Although vaccines and monoclonal antibody countermeasures have reduced the morbidity and mortality associated with SARS-CoV-2 infection, variants with constellations of mutations in the spike gene threaten their efficacy. Accordingly, antiviral interventions that are resistant to further virus evolution are needed. The host-derived cytokine IFN-λ has been proposed as a possible treatment based on correlative studies in human COVID-19 patients. Here, we show IFN-λ protects against SARS-CoV-2 B.1.351 (Beta) and B.1.1.529 (Omicron)variants in three strains of conventional and human ACE2 transgenic mice. Prophylaxis or therapy with nasally-delivered IFN-λ2 limited infection of historical or variant (B.1.351 and B.1.1.529) SARS-CoV-2 strains in the upper and lower respiratory tracts without causing excessive inflammation. In the lung, IFN-λ was produced preferentially in epithelial cells and acted on radio-resistant cells to protect against of SARS-CoV-2 infection. Thus, inhaled IFN-λ may have promise as a treatment for evolving SARS-CoV-2 variants that develop resistance to antibody-based countermeasures.

## INTRODUCTION

Severe acute respiratory syndrome coronavirus 2 (SARS-CoV-2) emerged in 2019 and has infected more than 300 million people worldwide. The coronavirus disease 2019 (COVID-19) pandemic continues because of the evolution of highly transmissible variant strains and a failure to vaccinate large segments of the global population. SARS-CoV-2 infection causes a range of influenza-like symptoms but can progress rapidly to pneumonia, acute respiratory distress syndrome (ARDS), and death (Guan et al., 2020; Huang et al., 2020).

One hallmark of COVID-19 in some individuals is a hyper-inflammatory state with excessive production of proinflammatory mediators, which recruit activated immune cells that ultimately impair alveolar gas-exchange and injure the lung (Mehta et al., 2020; Zhang et al., 2020). Interferons (IFNs) are pro-inflammatory cytokines that are a first line of defense against most virus infections. Type I (IFN-α subtypes and IFN-β) and type III IFNs (IFN-λs) are induced rapidly after detection by and activation of pathogen sensors (*e.g*., Toll-like [TLR] or RIG-I-like [RLR] receptors) and their downstream signaling pathways (Park and Iwasaki, 2020). Type I and III IFNs bind to distinct receptors on the cell surface to activate signal transducers and activators of transcription (STATs) proteins that induce expression of hundreds of antiviral IFN-stimulated genes (ISGs) (Lazear et al., 2019; Schneider et al., 2014). Cell culture studies have shown that IFN pre-treatment can restrict SARS-CoV-2 infection in human intestinal and airway epithelia (Felgenhauer et al., 2020; Stanifer et al., 2020; Vanderheiden et al., 2020). Although type I IFNs are a potential treatment strategy for SARS-CoV-2 infection (Hoagland et al., 2021), the ubiquitous expression of the IFNAR1/IFNAR2 receptor and strong, sustained pro-inflammatory responses can have pathological consequences. In comparison, the cellular response to type III IFN-λ is thought to be less inflammatory, as it functions primarily at epithelial and barrier surfaces where its heterodimeric receptor (IFNLR1/IL10Rβ) is preferentially expressed (Andreakos and Tsiodras, 2020; Broggi et al., 2020b; Galani et al., 2017).

The role of IFN-λ in SARS-CoV-2 infection and pathogenesis remains unclear. Although patients with severe COVID-19 patients have elevated serum levels of pro-inflammatory cytokines and chemokines, generally, their type I and III IFN levels are lower (Blanco-Melo et al., 2020; Galani et al., 2021), which suggests possible virus-induced antagonism or skewing of antiviral responses. Notwithstanding this point, in one human study, higher serum IFN-λ levels were associated with less viral infection in the respiratory tract and more rapid viral clearance, and a higher IFN-λ to type I IFN ratio correlated with improved outcome (Galani et al., 2021). In the respiratory tract, IFN-λ expression varies with location, level of viral burden, and degree of disease severity, and may have opposing roles at distinct anatomical sites in COVID-19 patients (Sposito et al., 2021). Thus, while IFN-λ expression appears to correlate inversely with COVID-19 severity, its mechanism(s) of protection is not well understood. Although IFN-λ has been studied in animals in the context of SARS-CoV-2 infection (Boudewijns et al., 2020; Broggi et al., 2020a; Dinnon et al., 2020; Sohn et al., 2021), and postulated to have a protective antiviral role, the responding cell types and targets of action have not been identified.

The emergence of SARS-CoV-2 variants (Beta, B.1.351; Gamma, B.1.1.28, Delta, B.1.617.2; and Omicron, B.1.1.529) with increasing antigenic divergence in the spike protein has highlighted a need for broad-spectrum antiviral agents that are less sensitive to viral evolution and the development of resistance. Hence, the potential benefits of host-target therapies, such as IFN-λ , have been discussed (Andreakos and Tsiodras, 2020; Prokunina-Olsson et al., 2020). Here, we investigated the potential efficacy of IFN-λ in the context of SARS-CoV-2 infection in mice. We found that *Ifnlr1^-/-^* (also termed IL28Rα^-/-^) C57BL/6 mice infected with B.1.351 or B.1.1.529 variants sustained higher viral burdens in the respiratory tract, indicating a protective role for IFN-λ against SARS-CoV-2 infection.

When we administered recombinant murine IFN-λ 2 by an intranasal route to K18-human (h)ACE2 transgenic mice or conventional 129S2 mice, as prophylaxis or therapy, we observed markedly reduced upper and lower respiratory tract infection and inflammation.

Administration of nasally-delivered IFN-λ 2 several days before or after infection conferred protection against infection in the lungs. IFN-λ was produced principally in epithelial cells and acted mainly on radio-resistant cells. Our data in mice suggest that IFN-λ has therapeutic potential as a less inflammatory, broad-spectrum antiviral agent against SARS-CoV-2 and its emerging variants.

## RESULTS

### IFN-λ signaling contributes to the antiviral response against SARS-CoV-2

To assess the importance of IFN-λ signaling in protection against SARS-CoV-2 infection, we inoculated 6-week-old wild-type (WT) and congenic *Ifnlr1^-/-^* C57BL/6 mice with 10^5^ focus-forming units (FFU) of SARS-CoV-2 B.1.351 virus, which contains K417Y, E484K, and N501Y substitutions in the spike receptor-binding domain (RBD) (Tegally et al., 2021). Prior studies have shown that the N501Y change in spike is mouse-adapting and can enable binding to mouse ACE2 and infection of several laboratory strains of mice (Chen et al., 2021a; Li et al., 2021; Rathnasinghe et al., 2021; Shuai et al., 2021; Winkler et al., 2021; Zhang et al., 2021a). *Ifnlr1^-/-^* mice showed higher viral RNA levels at 7 days post infection (dpi) in nasal washes and lung homogenates compared to WT mice (**Fig 1A**). Consistent with these data, we detected substantially higher levels of infectious virus by plaque assay in the

**Figure 1.**
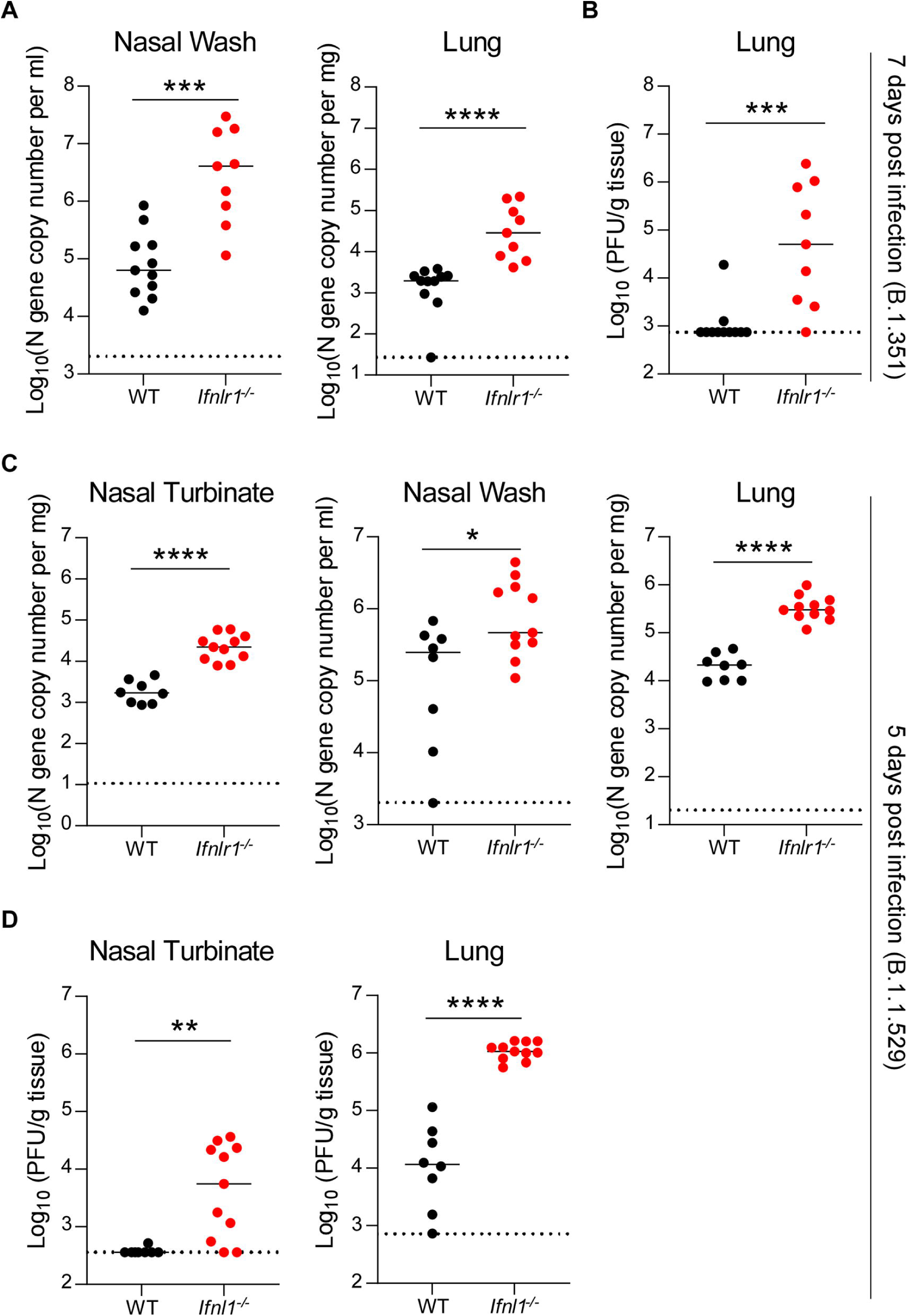
Increased susceptibility of *Ifnlr1*^-/-^ mice to SARS-CoV-2 infection. (**A-B**) Six-week-old male and female C57BL/6 WT and *Ifnlr1*^-/-^ mice were inoculated with 10^5^ FFU of B.1.351. (**A**) Viral RNA levels were measured from tissues at 7 dpi by qRT-PCR. (**B**) Infectious virus was measured from tissues by plaque assay at 7 dpi (n = 9-11 per group, 2 experiments). (**C-D**) Three-month-old female and male C57BL/6 WT and *Ifnlr1*^-/-^ mice were inoculated with 10^5^ FFU of B.1.1.529. (**C**) Viral RNA levels were measured at 5 dpi by qRT-PCR. Note, an earlier time point of analysis was used because B.1.1.529 is less pathogenic in mice. (**D**) Infectious virus was measured by plaque assay at 5 dpi (n = 8-11 per group, 2 experiments). Bars indicate median values. Data were analyzed by Mann-Whitney test (\**P* < 0.05, \*\**P* < 0.01, \*\*\**P* < 0.001, and \*\*\*\**P* < 0.0001).

lungs of *Ifnlr1^-/-^* mice at 7 dpi (**Fig 1B**). Next, we investigated whether IFN-λ also had protective effects against the emerging SARS-CoV-2 B.1.1.529 Omicron variant, which has mutations that could enable evasion against vaccines and therapeutic antibodies (Zhang et al., 2021b), We inoculated 3-month-old WT and *Ifnlr1^-/-^* mice with 10^5^ FFU of B.1.1.529 and observed that *Ifnlr1^-/-^* mice sustained higher levels of viral RNA in nasal turbinates, nasal washes, and lungs at 5 dpi (**Fig 1C**). Infectious virus titers also were higher in *Ifnlr1^-/-^* than WT mice in both nasal turbinates and lung homogenates (**Fig 1D**). Collectively, these data suggest that IFN-λ signaling has an antiviral role during SARS-CoV-2 variant infection in C57BL/6 mice.

### Exogenous IFN-**λ**2 limits SARS-CoV-2 virus infection and inflammation in **K18-hACE2 transgenic mice.**

We next evaluated the protective activity of exogenous IFN-λ 2 against SARS-CoV-2 infection in mice. In a first set of experiments, we used K18-hACE2 transgenic mice, which express hACE2 under regulation of the epithelial cell cytokeratin-18 promoter and are highly vulnerable to SARS-CoV-2-induced pneumonia and brain infection (Golden et al., 2020; Oladunni et al., 2020; Winkler et al., 2020). We first administered 2 μg of commercially-available IFN-λ 2 via intranasal or intraperitoneal route 16 h before inoculation with a historical WA1/2020 D614G SARS-CoV-2 strain. At 3 dpi, mice treated with IFN-λ 2 by an intranasal route had markedly lower levels of viral RNA and infectious virus in the nasal turbinates, nasal washes, lungs, and brain (**Fig S1B-C**), whereas animals treated by an intraperitoneal route did not show these reductions (**Fig S1A**). Based on these data, we used intranasal administration of IFN-λ 2 for the remainder of our studies. We extended the window of prophylaxis in K18-hACE2 mice with a single intranasal dose of IFN-λ 2 at day -2 (D-2) or -3 (D-3) before inoculation with WA1/2020 D614G. IFN-λ 2 treatment at D-2 resulted in lower viral RNA levels in nasal turbinates, nasal washes, and lungs, but not in the brain at 3 dpi (**Fig 2A** **and S2A**). Infectious virus levels in the lungs of IFN-λ 2-treated animals were lower than in PBS-treated animals; however, there was no difference in the nasal turbinates of these two groups (**Fig 2B**). D-3 treatment with IFN-λ 2 showed reduced viral RNA and infectious virus levels in the lungs at 3 dpi but not in other tissues (**Fig S1D-E**). Finally, we tested whether protection could be improved with two doses of IFN-λ 2 treatment, one administered before and a second given after virus inoculation. K18-hACE2 mice were treated with 2 μg of IFN-λ2 via intranasal route at 16 h before and 8 h after intranasal inoculation with 10^3^ FFU of WA1/2020 D614G. Notably, IFN-λ2 treatment prevented weight loss (**Fig S1H**) and showed reduced levels of viral RNA and infectious virus at 7 dpi in the nasal turbinates, nasal washes, lungs and brain compared to PBS-treated mice (**Fig S1I-J**).

We next explored the therapeutic efficacy of IFN-λ 2. K18-hACE2 mice were administered a single 2 μg dose of IFN-λ 2 via nasal route at 8 h after infection, and animals were sacrificed at 3 dpi. IFN-λ 2 treated mice showed reduced viral RNA levels in the nasal turbinates, lungs, and brain (**Fig S1F**), and infectious virus titers in the nasal turbinates and lungs (**Fig S1G**). However, therapeutic administration of IFN-λ 2 did not reduce viral burden in nasal washes compared to PBS-treated animals (**Fig S1F**). We also administered IFN-λ 2 as a two-dose therapy at 1 (D+1) and 2 (D+2) dpi, which resulted in lower viral RNA loads in nasal turbinates and lungs, but not in nasal washes or the brain (**Fig 2C** **and S2B**). Infectious virus levels also were lower in the lungs with this IFN-λ 2 treatment scheme (**Fig 2D**).

**Figure 2.**
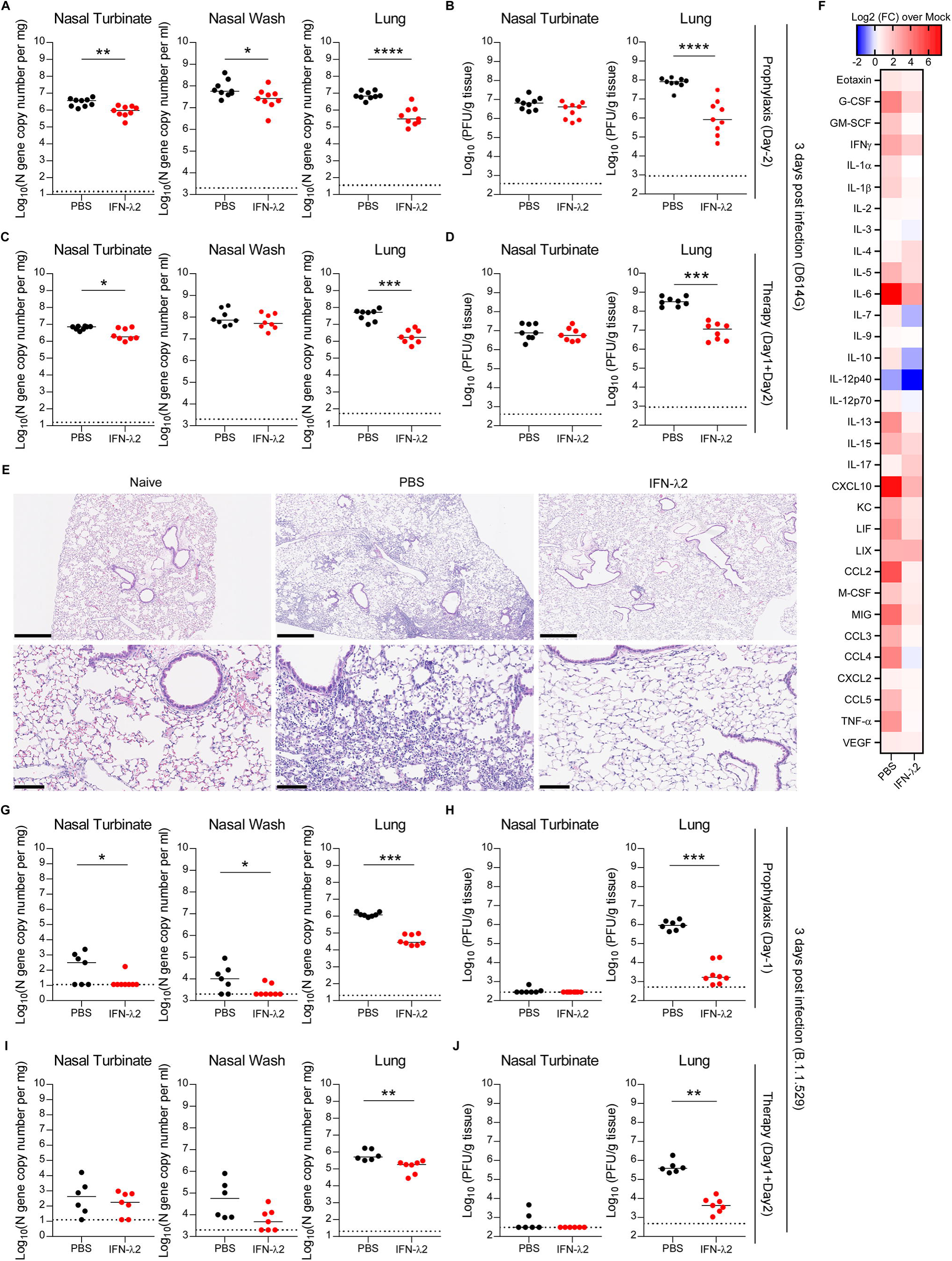
Nasally-delivered IFN-λ 2 treatment protects K18-hACE2 mice against SARS-CoV-2 infection. Eight-week-old female K18-hACE2 mice were inoculated by intranasal route with 10^3^ FFU of WA1/2020 D614G. At D-2 (**A-B**) or D+1 and D+2 (**C-D**), mice were given a single 2 g dose of murine IFN-λ 2 or PBS by the intranasal route. (**A and C**) Viral RNA levels were measured at 3 dpi. (**B and D**) Infectious virus was measured at 3 dpi (**A-B**: n = 9 per group, 2 experiments; **C-D**: n = 8 per group, 2 experiments). (**E**) Hematoxylin and eosin staining of lung sections from animals treated with 2 μg doses of murine IFN-λ 2 or PBS by intranasal route at -16 h and +8 h relative to inoculation with WA1/2020 D614G and harvested at 7 dpi. Low (top, scale bars, 500 μm) and high (bottom, scale bars, 100 μm) power images are shown. Representative images from n = 5 per group. (**F**) Eight-week-old female K18-hACE2 mice were treated with 2 μg of murine IFN-λ2 or PBS at -16 h and challenged with 10^3^ FFU of WA1/2020 D614G. Heat-maps of cytokine levels in lung homogenates at 3 dpi. Fold-change was calculated relative to mock-infected mice, and log_2_ values are plotted (2 experiments, n = 7 per group except naïve, n = 4). (**G-J**) Five-month-old female K18-hACE2 mice were inoculated with 10^3^ FFU of B.1.1529. At D-1 (**G-H**) or D+1 and D+2 (**I-J**), mice were given 2 g of murine IFN-λ 2 or PBS by the intranasal route. Viral RNA (**G and I**) and infectious (**H and J**) virus levels were measured at 3 dpi (**G-H**: n = 7-8 per group, 2 experiments; **I-J**: n = 6-7 per group, 2 experiments). Bars (**A-D** and **G-J**) indicate median values. Data were analyzed by Mann-Whitney tests (**A-D** and **G-J**) (\**P* < 0.05, \*\**P* < 0.01, \*\*\**P* < 0.001, and \*\*\*\**P* < 0.0001).

Some COVID-19 patients develop hyper-inflammatory immune responses, which may contribute to respiratory failure (Andreakos and Tsiodras, 2020; Galani et al., 2021). Given that IFN-λ 2 treatment reduced viral levels in the lung, we hypothesized that it also might mitigate immune responses and lung disease. Lung tissues were collected from IFN-λ 2 or PBS-treated mice at 7 dpi and sectioned for histological analysis; this time point was selected since lung pathology in K18-hACE2 mice is greater at 7 than 3 dpi. Lungs from PBS-treated, SARS-CoV-2-infected K18-hACE2 mice showed diffusely infiltrating immune cells with alveolar space consolidation consistent with pneumonia, whereas this was observed to a substantially lesser degree in IFN-λ2-treated animals (**Fig 2E**). Measurement of cytokine and chemokines in lung homogenates at 3 dpi showed decreased levels of G-CSF, IL-1β, IL-6, CXCL10, CCL2, and TNF-αin IFN-λ 2-treated K18-hACE2 mice (**Fig 2F** **and S3**). These results suggest that treatment with IFN-λ2 can protect mice against SARS-CoV-2 by inhibiting lung infection and inflammation.

We evaluated whether exogenous IFN-λ 2 treatment could also protect K18-hACE2 mice from the B.1.1.529 Omicron variant. First, we administered mice a single 2 μg dose of IFN-λ 2 at D-1. IFN-λ 2 treated mice had lower levels of B.1.1.529 viral RNA in nasal turbinates, nasal washes and lungs (**Fig 2G**) as well as infectious virus in lungs than PBS-treated animals (**Fig 2H**). Our two-dose therapeutic regimen at D+1 and D+2 also reduced levels of B.1.1.529 viral RNA and infectious virus in the lungs, but not in the nasal turbinates or washes (**Fig 2I-J**). While performing these studies, we observed an absence of viral RNA in the brain of PBS-treated B.1.1.529-infected K18-hACE2 mice (**Fig S2C-D**) and low levels of infection in nasal turbinates (**Fig 2H and J**), which is consistent with recent studies suggesting B.1.1.529 is less pathogenic in rodents (Diamond et al., 2021).

Nonetheless, our experiments demonstrate that exogenous IFN-λ 2 protects against B.1.1.529 infection in K18-hACE2 mice.

### Exogenous IFN-λ 2 limits SARS-CoV-2 infection and inflammation in 129S2 mice

To confirm our results in another model of SARS-CoV-2 infection, we treated and challenged 129S2 mice, which are susceptible to SARS-CoV-2 strains (*e.g*., B.1.351) with an N501Y mouse-adapting mutation, more so than C57BL/6 mice (Chen et al., 2021a; Li et al., 2021; Rathnasinghe et al., 2021; Shuai et al., 2021; Zhang et al., 2021a). Nasal administration of IFN-λ 2 at D-1 protected B.1.351-infected mice from weight loss (**Fig 3A**) and reduced viral burden in nasal turbinates, nasal washes, lungs, and brain (**Fig 3B-C** **and S2E**). When we extended the prophylaxis window to D-3 or D-5, IFN-λ 2 still reduced infection-induced weight loss (**Fig 3D and G**) and viral RNA and infectious virus levels in nasal turbinates and lungs, but not in nasal washes (**Fig 3E, F, H and I**). 129S2 mice treated at D-3 but not D-5 with IFN-λ 2 had less viral RNA in the brain that those administered PBS (**Fig S2F-G**). We next evaluated the effect of two 2- μg doses of IFN-λ 2 -16 h and +8 h infection on B.1.351 infection. Infected 129S2 mice treated with PBS showed about 15% weight loss by 4 dpi, whereas IFN-λ2 treated animals did not (**Fig 3J**). Levels of viral RNA and infectious virus levels were reduced in the nasal turbinates, nasal washes, lungs, and brain of IFN-λ2-treated compared to PBS-treated mice (**Fig 3K-L** **and S2H**).

**Figure 3.**
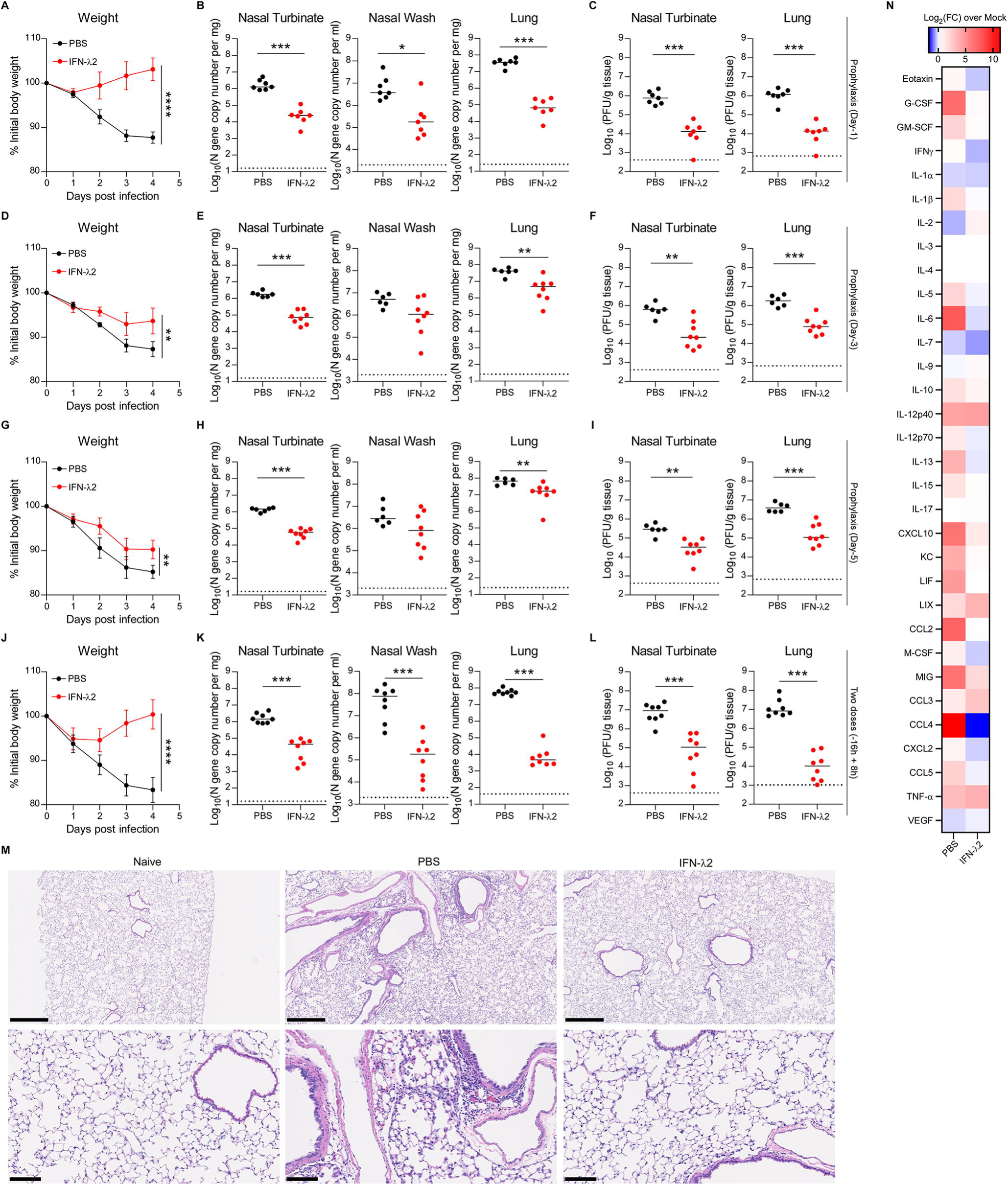
IFN-λ 2 treatment protects 129S2 mice against SARS-CoV-2 infection. **(A-I)** Six-week-old female 129S2 mice were inoculated by intranasal route with 10^5^ FFU of B.1.351. At D-1 (**A-C**), D-3 (**D-F**) or D-5 (**G-I**), mice were given a single 2 μ murine IFN-λ 2 or PBS by intranasal route. (**A, D, G**) Weight change. (**B, E, H**) Viral RNA levels at 4 dpi. (**C, F, I**) Infectious virus levels at 4 dpi (**A-C**: n = 7 per group, 2 experiments; **D-F**: n = 6-8 per group, 2 experiments; **G-I**: n = 6-8 per group, 2 experiments). (**J-L**) Six-week-old female 129S2 mice were inoculated by intranasal route with 10^5^ FFU of B.1.351. At -16 h and +8 h, mice were administered 2 g of murine IFN-λ 2 or PBS by intranasal route. (**J**) Weight change. (**K**) Viral RNA levels at 4 dpi. (**L**) Infectious virus levels at 4 dpi in (n = 8 per group, 2 experiments). (**M**) Hematoxylin and eosin staining of lung sections at 4 dpi from animals treated in (**J-L**). Low (top, scale bars, 500 μm) and high (bottom, scale bars, 100 μm) power images are shown (representative of n = 5 per group). (**N**) Heat-maps of cytokine levels in lung homogenates at 4 dpi from animals treated in (**J-L**). Fold-change was calculated compared to mock infected mice, and log_2_ values were plotted (n = 8 per group except naïve n = 4, 2 experiments). Bars (**B-C, E-F, H-I** and **K-L**) indicate median values. Data were analyzed by Mann-Whitney tests (**B-C, E-F, H-I** and **K-L**) or *t* tests of the area under the curve (**A, D, G and J**) (\**P* < 0.05, \*\**P* < 0.01, \*\*\**P* < 0.001, and \*\*\*\**P* < 0.0001).

Lung sections from B.1.351-infected, PBS-treated 129S2 mice at 4 dpi showed mild to moderate immune cell infiltration, extravasation of erythrocytes into the alveolar space, and pulmonary vascular congestion, whereas those treated with IFN-λ2 appeared more like uninfected, naive mice (**Fig 3M**). Consistent with these data, IFN-λ2 treated mice had reduced levels of the pro-inflammatory cytokines and chemokines that were elevated in B.1.351-infected PBS treated mice including IL-1β, IL-6, CXCL10, CCL2, CCL4, and CCL5 (**Fig 3N** **and S4**). Collectively, our data establishes a protective effect of IFN-λ2 against SARS-CoV-2 infection in multiple strains of mice.

### IFN-λ2 transcriptional signature in the lung

To begin to understand how IFN-λ2 protects against SARS-CoV-2 in the lung, we performed bulk RNA sequencing on tissues obtained from naïve animals or animals treated IFN-λ2 via the intranasal route. Principal component analysis showed distinct transcriptional signatures in the lungs of IFN-λ2-treated mice at 1 (D+1) or 3 (D+3) day(s) after treatment compared to naïve mice. The transcriptional signature in the lung at D+1 after IFN-λ2 was distinct from naïve animals, whereas by D+3 the signature started to return to baseline (**Fig 4A**). We identified 1,820 and 1,317 differentially expressed genes (DEGs) in the D+1 and D+3 IFN-λ2-treated groups, respectively, and 856 DEGs were identified between the D+1 and D+3 groups (**Fig 4B**). We performed Metascape analysis to define biological pathways enriched in the IFN-λ2-treated groups compared to naïve group. Among the top enriched up-regulated pathways in both the D+1 and D+3 groups relative to the naïve group were extracellular matrix organization signaling (*e.g*., *Col2a1*, *Col5a2*, *Lampb3,* and *Mmp15*), regulation of cell adhesion signaling (*e.g*., *Vegfc*, *Jam2,* and *Cav1*), response to wounding signaling (*e.g*., *CD36*, *Timp1,* and *Col3a1*), and negative regulation of cytokine production signaling (*e.g*., *Klf2*, *Arg2,* and *Foxj1*) (**Fig 4C-D** **and S5**). Although these pathways were enriched in both groups, expression of these genes in D+3 group was lower (**Fig 4C-D** **and S5**), suggesting the effect of IFN-λ 2 had begun to wane. In comparison, other transcriptional programs were uniquely expressed in D+1 group including response to IFN-α signaling (*e.g*., *Oas1a*, *Ifit2,* and *Bsl2*) and virus signaling (*e.g*., *Cxcl10*, *Rsad2*, *Isg15*, *Irf7,* and *Ifit1*) (**Fig 4C-D**), suggesting these antiviral signals are induced quickly and decline rapidly once the stimulus is lost. Other pathways transcriptionally induced by IFN-λ2 at D+1 only included T cell mediated cytotoxicity signaling (*e.g*., *H2-q1*, *H2-q7*, *H2-k1,* and *Tap2*) and morphogenesis of a branching epithelium signaling (*e.g*., *Wnt2*, *Foxc2,* and *Myc*) (**Fig 4C-D** **and S5**). Biological pathways that were downregulated in D+1 and D+3 groups compared to naïve samples included sodium ion transport signaling, protein citrullination signaling and potassium ion transmembrane transport signaling. Some pathways that were downregulated only in the D+1 group included responses to xenobiotic stimulus signaling and negative regulation of lipid metabolic process signaling (e.g., *Apobec1*, *Serpina12* and *Gper1*).

**Figure 4.**
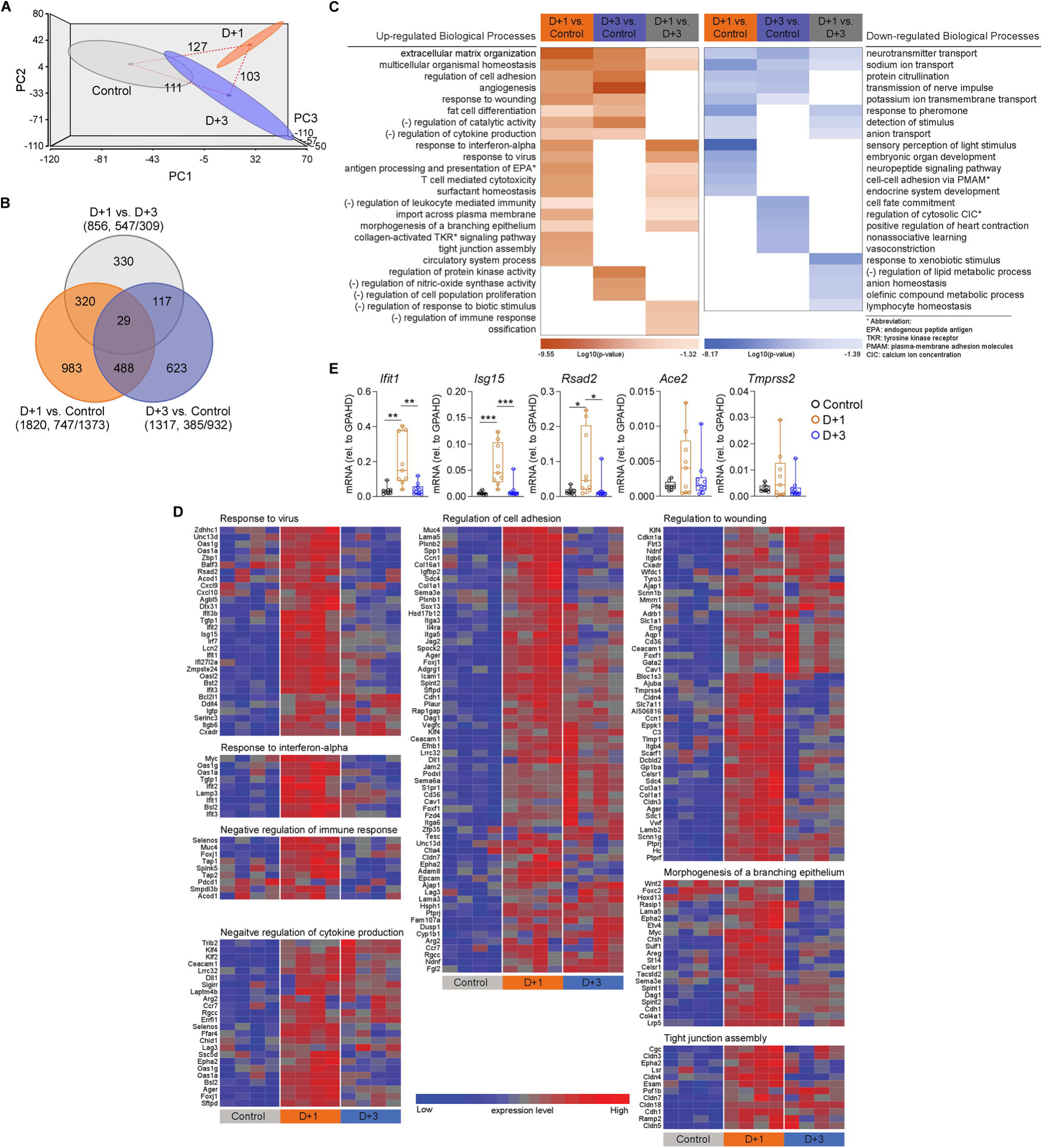
Transcriptional signatures in the lungs of mice treated with murine IFN-λ2. (**A-D**) RNA sequencing of lung homogenates of naive female K18-hACE2 mice (control) or mice treated with 2 g of murine IFN-λ 2 by intranasal route for 1 (Day+1) or 3 (Day+3) days. (**A**) Three-dimensional map from principal component analysis. Each group is represented by an ellipse and the color-matched solid circle, which is the centroid of each group. The size of the ellipse is the centroid with one standard deviation. The dashed red lines with numbers indicate the spatial distance between centroids of the 3 groups, which is calculated by using the three-dimensional coordinates for the centroids. (**B**) Venn diagrams of overlapping genes identified in differential expression analysis when comparing with control, D+1, and D+3 groups. Numbers in the parenthesis under each comparison indicate differentially expressed genes (fold-change ≥ 2 at *P* < 0.05) followed by the proportion that are up- or downregulated. (**C**) The significantly enriched biological processes defined by a Metascape pathway analysis tool comparing control, D+1, and D+3 groups; upregulated genes (brown) or downregulated (blue) in the IFN- λ 2 treated group (D+1 or D+3) compared to the control group or in the D+1 group compared to the D+3 group. (**D**) Heatmaps of selected biological processes enriched in the D+1 group or the D+3 group versus the control group (n = 4 per group). (**E**) mRNA levels of indicated target genes were measured from the lung homogenates of naive female K18-hACE2 mice or mice treated with 2 μg of murine IFN-λ 2 by intranasal route for D+1 or D+3 days (n = 8-10 per group, 2 experiments). Data in (**E**) were analyzed by one-way ANOVA with Dunnett’s post-test (\**P* < 0.05, \*\**P* < 0.01, and \*\*\**P* < 0.001).

We validated our bulk RNA sequencing data by qRT-PCR by measuring expression of several ISGs including *Ifit1*, *Isg15,* and *Rsad2* that can respond to IFN-λ signaling (Jilg et al., 2014; Lazear et al., 2019; Shindo et al., 2013). Notably, these ISGs expression levels were upregulated at D+1 and diminished at D+3 (**Fig 4E**). We did not observe changes in mRNA expression of *Ace2*, which can be modulated by type I IFN (Ziegler et al., 2020), or *Tmprss2* (**Fig 4E**), two key genes involved in SARS-CoV-2 attachment and entry, suggesting they do not respond to IFN-λ signals in mice. Collectively, our data demonstrate that the transcriptional program induced by IFN-λ2 is characterized by a short burst of expression of antiviral, cell-to-cell communication, and wound healing gene programs. However, we did not observe higher levels of NF-κB genes (*e.g*., *Il6, Il1*β and *Tnf*α), which can be strongly induced by type I IFN (Galani et al., 2017), indicating IFN-λ selectively induces antiviral but not highly pro-inflammatory genes.

### IFN-λ is preferentially produced by epithelial cells during SARS-CoV-2 infection

We investigated which cell type(s) in the lung produce IFN-λ after SARS-CoV-2 infection *in vivo*. *Ifnl2* and *Ifnl3* mRNA expression levels was upregulated at 2 dpi after WT C57BL/6 mice were inoculated with 10^6^ FFU of B.1.351 (**Fig 5A**). To identify the cell types expressing IFN-λ mRNA, at 2 dpi we sorted under BSL3 conditions lung epithelial cells (ECs) and different immune cells populations (alveolar macrophages (AM), monocytes (Mo), neutrophils (Nϕ), B cells (B), T cells (T), and dendritic cells (DC)) and then performed qRT-PCR for the two IFN-λ transcripts in mice (**Fig 5B** **and S6**). CD45^-^CD326^+^ lung ECs had the highest levels of *Ifnl2* and *Ifnl3* mRNA expression with CD45^+^CD11c^+^ Siglec F^-^MHCII^+^ DCs showing the next highest expression; the other cell types analyzed had limited mRNA expression of *Ifnl2* and *Ifnl3* (**Fig 5C**). As expected, based on the literature (Galani et al., 2017; Lazear et al., 2019), the *Ifnlr1* receptor was expressed mainly on CD45^-^CD326^+^ ECs and CD45^+^CD11b^+^Ly6G^+^ Nϕ (**Fig 5C**). To corroborate these results, we utilized *Ifnl2-Egfp* reporter mice (Galani et al., 2017) to evaluate IFN-λ expression. EGFP was greatly induced at 2 dpi and localized mostly to CD326^+^ ECs lining the bronchial walls; however, we did not observe substantial EGFP signal in the lung parenchyma (**Fig 5D**). We also investigated the tropism of SARS-CoV-2 B.1.351 after infection in the lung.

**Figure 5.**
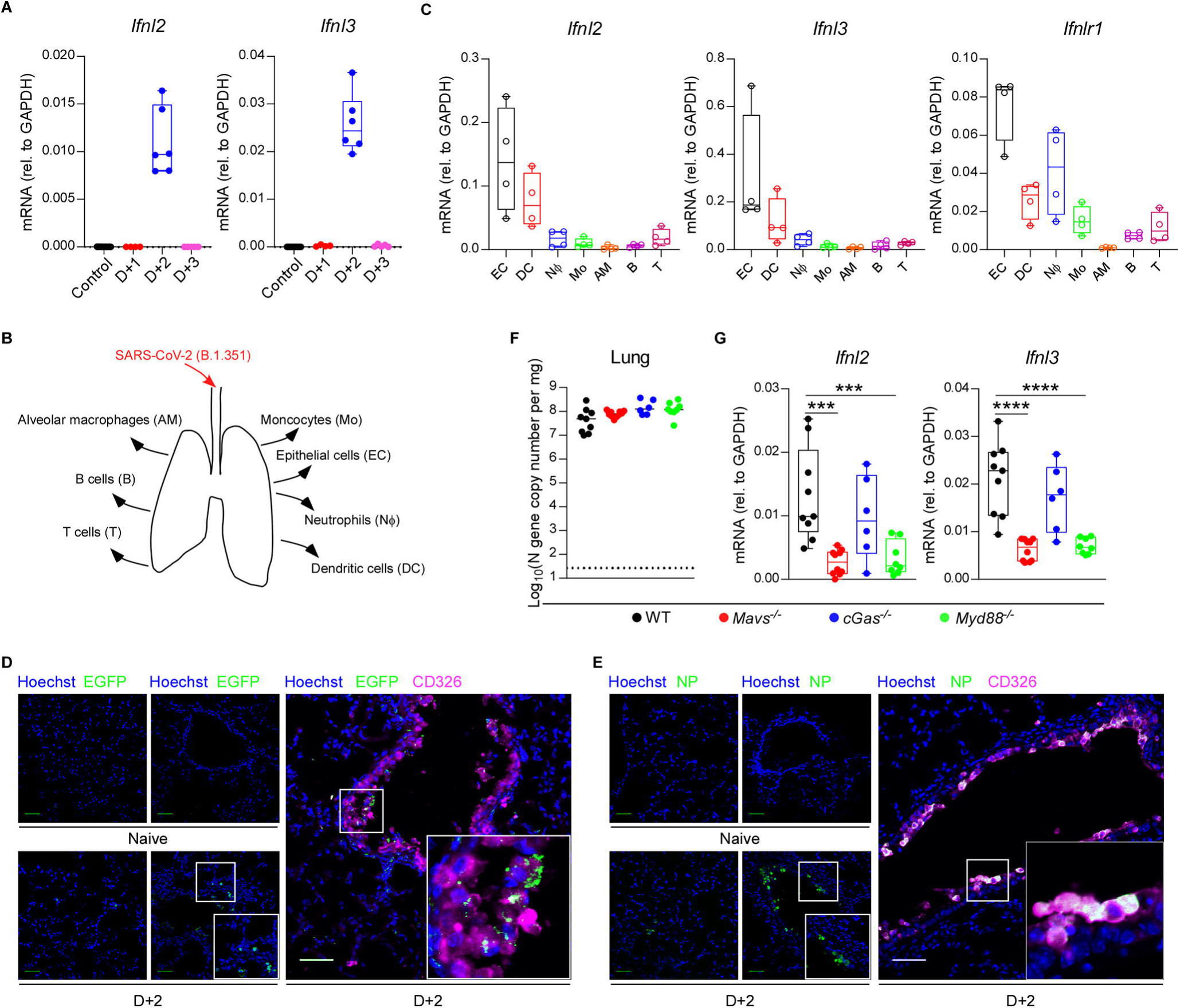
IFN-λ expression *in vivo*. (**A**) Six-week-old male and female C57BL/6 mice were inoculated with 10^6^ FFU of B.1.351. *Ifnl2* and *Ifnl3* mRNA levels from lungs were measured at indicated days post infection by qRT-PCR (n = 4-9 per group, 2 experiments). (**B-C**) Six-week-old male and female C57BL/6 mice were inoculated with 10^6^ FFU of B.1.351. (**B**) Scheme depicting cell populations isolated from lungs of B.1.351-infected mice. (**C**) *Ifnl2*, *Ifnl3* and *Ifnlr1* mRNA expression levels from (**B**) were measured by qRT-PCR at 2 dpi (n = 4 per group, each dot represents 4 mice pooled together, 2 experiments). (**D-E**) Six-week-old C57BL/6 or *Ifnl2-Egfp* reporter mice were inoculated with 10^6^ FFU of SARS-CoV-2 B.1.351. (**D**) Localization of EGFP and epithelial cells (ECs, CD326) in the lungs of *Ifnl2-Egfp* reporter mice at 2 dpi. Frozen sections stained for GFP (green), CD326 (magenta), and Hoechst (blue) are shown. Scale bar: 50 μm. (**E**) Localization of SARS-CoV-2 positive and ECs in the lung of mice at 2 dpi. Frozen sections stained for SARS-CoV-2 nucleocapsid protein (NP) (green), CD326 (magenta), and Hoechst (blue) are shown. Scale bars in (**D-E**): 50 μm. (**F-G**) Six-week-old male and female WT, *Mavs* , *cGas* , or *Myd88^-/-^* C57BL/6 mice were inoculated with 10^6^ FFU of B.1.351. Viral RNA levels (**F**) or *Ifnl2* and *Ifnl3* mRNA expression levels (**G**) from lungs were measured at 2 dpi by qRT-PCR (n = 6-10, 2 experiments). Bars in (**F**) indicate median values. Data in (**F-G**) were analyzed by one-way ANOVA with Dunnett’s post-test (\*\*\**P* < 0.001 and \*\*\*\**P* < 0.0001).

Immunofluorescence microscopy for SARS-CoV-2 nucleocapsid showed that virus localized to airway tract epithelium and co-localized with CD326^+^ ECs (**Fig 5E**). This pattern suggests that ECs are the dominant cell type targeted for infection by SARS-CoV-2 and major source of IFN-λ production in the lower respiratory tract We next evaluated which pathogen recognition receptor signaling pathways induced IFN-λ expression. Since IFNs can be activated though TLRs, RLRs, or cGAS-STING pathways after viral infections (Park and Iwasaki, 2020), we repeated B.1.351 infection experiments in *Mavs*^-/-^, *cGas*^-/-^ and *Myd88*^-/-^ C57BL/6 mice. At 2 dpi, levels of *Ifnl2* and *Ifnl3* mRNA in the lung were remarkably decreased in both *Mavs*^-/-^ and *Myd88^-/-^* mice, but not in *cGas*^-/-^ mice compared to WT mice (**Fig 5F**). Viral RNA levels were relatively equivalent among different mouse genotypes at this early time point (**Fig 5E**), suggesting the differences in IFN-λ expression levels were not skewed by effects on viral burden, and that the antiviral effect conferred by IFN-λ in the lung requires several days to manifest. Overall, our data suggest that in the lungs of mice after SARS-CoV-2 infection, IFN-λ is produced principally by epithelial cells though both MAVS and MyD88-dependent signaling pathways.

### IFN-λ signaling in radio-resistant cells controls SARS-CoV-2 infection in the lung

As our RT-PCR data demonstrated, in the lung, IFN-λ receptors (IFNLR1/IL10Rβ) are expressed in epithelial cells and some immune cells including neutrophils (Broggi et al., 2017; Lazear et al., 2019). To determine which cell type contributed to the protective effect mediated by IFN-λ against SARS-CoV-2 *in vivo*, we first depleted circulating neutrophils with anti-Ly6G [1A8 mAb] in the context of IFN-λ2 treatment (**Fig S7A**). Depletion of neutrophils had no impact on the reduction in weight loss or viral burden conferred by IFN-λ2 (**Fig 6A-C**). We next generated reciprocal sets of chimeric animals in which the radio-resistant compartment or radio-sensitive hematopoietic cells lacked that capacity for IFN-λ signaling using donor WT or *Inflr1*^-/-^ bone narrow and sublethally irradiated WT or *Inflr1*^-/-^ recipient mice (**Fig 6D** **and S7B**). Animals lacking IFN-λ signaling in the radio-resistant compartment sustained similar levels of infection in the nasal washes as fully *Ifnlr1^-/-^* mice, whereas animals lacking IFN-λ signaling in hematopoietic cells had similar levels of viral RNA as mice with intact IFN-λ signaling in all cells (**Fig 6E**). In the lungs, the similar trends were observed with higher levels of viral RNA in animals lacking *Ifnlr1* in the radio-resistant cell compartment (**Fig 6E**). In the nasal turbinates, the data was more nuanced, where both radio-resistant and radio-sensitive *Ifnlr1* signaling cell populations appear to contribute to IFN-λ-dependent control of SARS-CoV-2 infection (**Fig 6E**). Overall, our data suggest IFN-λ signaling protects mice against SARS-CoV-2 infection and depends dominantly on signaling in radio-resistant cells in the lung.

**Figure 6.**
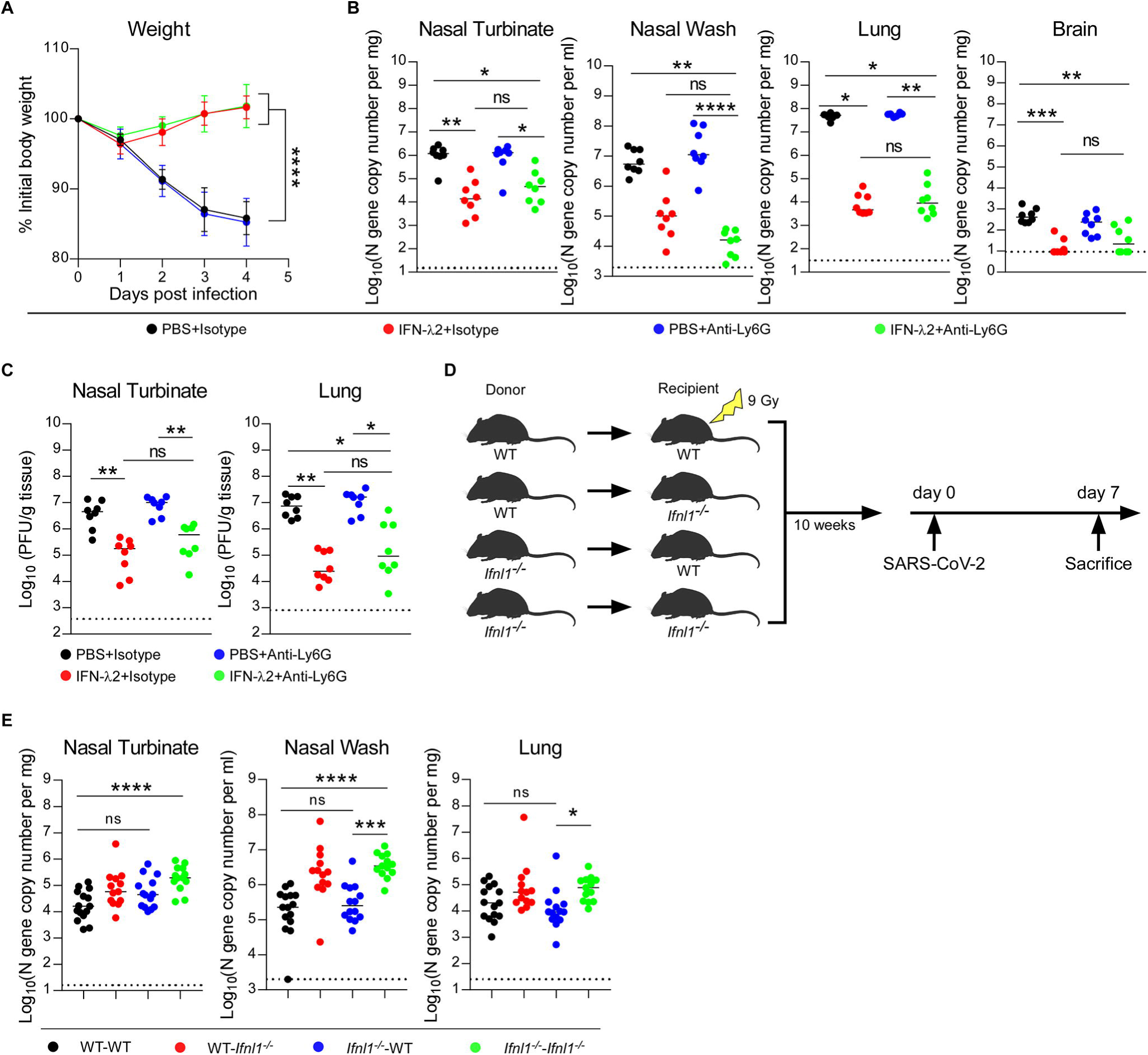
IFN-λ signaling in radio-resistant cells protects against SARS-CoV-2 infection. (**A-C**) Six-week-old female 129S2 mice received anti-Ly6G [1A8] or isotype control antibodies by intraperitoneal injection at D-1, D+1 and D+3 relative to B.1.351 infection (10^5^ FFU). Mice also were treated with 2 μg of murine IFN-λ2 or PBS at -16 h and +8 h by the intranasal route. (**A**) Weight change. (**B**) Viral RNA levels at 4 dpi. (**C**) Infectious virus levels at 4 dpi (n = 8 per group, 2 experiments). (**D**) Experimental scheme for generating of WT and *Ifnlr1^-/-^* bone marrow chimeric mice. Ten weeks after irradiation, mice were inoculated by the intranasal route with 10^5^ FFU of B.1.351. (**E**) Viral RNA levels at 7 dpi (n = 13-15 per group, 3 experiments). Bars indicate median values. Data were analyzed by one-way ANOVA with Dunnett’s post-test of the area under the curve (**A**) and Kruskal-Wallis test with Dunnett’s post-test (**B**-**C** and **E**) (\**P* < 0.05, \*\**P* < 0.01, \*\*\**P* < 0.001, and \*\*\*\**P* < 0.0001).

## DISCUSSION

In humans and other animals, SARS-CoV-2 targets the respiratory tract, which can result in the development of pneumonia, ARDS, and death (Guan et al., 2020; Huang et al., 2020). While existing neutralizing antibodies and vaccines against SARS-CoV-2 have conferred protection for many individuals, their efficacy is jeopardized by emerging variants that have increasing numbers of amino acid substitutions in the spike protein (Baum et al., 2020; Chen et al., 2021b; Hoffmann et al., 2021; Liu et al., 2021). Thus, therapeutic approaches are needed that can overcome viral resistance. IFN-λ induces hundreds of ISGs and has protective functions against many different virus infections, at least in cell culture and animal models (Lazear et al., 2019; Park and Iwasaki, 2020). Also, IFN-λ preferentially functions at mucosal sites including the respiratory tract because of the selected cellular expression of IFNLR1, a subunit of its receptor (Broggi et al., 2020b; Lazear et al., 2019). While type I IFN is also antiviral and has greater potency, treatment is often associated with collateral systemic effects and inflammation. For these reasons, we investigated the potential of IFN-λ in preventing and treating SARS-CoV-2 infection. Our data in mice show that IFN-λ can protect against infection by two variants (B.1.351 and B.1.1.529) and diminish inflammatory responses in the lung. In the context of SARS-CoV-2 infection, IFN-λ in the lung was produced primarily by ECs and acted on radio-resistant cells to confer protection.

Host-derived innate immune responses have the potential to limit the impact of viral evolution since multiple genes and pathways contribute to inhibitory responses. Nonetheless, virus-mediated attenuation of innate immune antiviral response occurs and is linked to SARS-CoV-2 disease severity (Blanco-Melo et al., 2020; Galani et al., 2021; Sposito et al., 2021). Indeed, serum IFN-λ levels are low in patients with severe COVID-19, yet those with higher levels have better outcomes (Blanco-Melo et al., 2020; Galani et al., 2021). Related to this, high levels of IFN-λ in the upper respiratory tract were associated with higher viral burden but less disease severity, whereas patients with severe COVID-19 had elevated IFN-λ levels in the lower respiratory tract (Sposito et al., 2021). In mice, we detected IFN-λ gene expression in the lung within days of SARS-CoV-2 infection, and *Ifnlr1^-/-^* mice lacking IFN-λ signaling sustained higher viral burden in the upper and lower respiratory tract, suggesting IFN-λ can protect against SARS-CoV-2 infection *in vivo*.

Because of the potential of IFN-λ as a broadly-acting therapy, we evaluated its antiviral activity *in vivo*. Notably, equivalent doses of IFN- λ2 delivered by a nasal but not systemic route could limit SARS-CoV-2 infection. The basis for this disparity remains uncertain, although higher doses given by a peripheral route might have protective effects, as was seen by others after subcutaneous administration of pegylated forms of IFN-λ (Dinnon et al., 2020). Post-exposure therapy with IFN-λ2 also conferred protection in the lung in mice, but the antiviral effects in other tissues were diminished, suggesting that once infection is established in the upper airway, and viral evasion mechanisms are induced, IFN-λ2 therapy may have less benefit. By testing several key variants (B.1.351 and B.1.1.1529), we established that IFN-λ2 could protect broadly against antigenically-distinct SARS-CoV-2 isolates, and thus may be less susceptible to immune escape than monoclonal or serum-derived antibodies (Baum et al., 2020; Chen et al., 2021a; Hoffmann et al., 2021; Liu et al., 2021).

Even a single dose of IFN-λ at D-5 conferred mice protection, demonstrating a persistent antiviral effect. The basis for this durable inhibitory effect remains uncertain especially in light of our transcriptional profiling data in the lung, which showed a rapid induction and then dampening of gene induction. Although further studies are warranted to define the basis of the durable inhibitory effect, we speculate that the half-life of certain inhibitory ISG products may be longer, transcriptional activation downstream of IFN-λ signaling may have distinct kinetics in the upper airway, or immune cells may become ‘trained’ (Netea et al., 2020) by IFN-λ to respond more quickly. We used soluble IFN-λ in our administration scheme. It remains possible that the window of prevention and clinical utility could be extended further by administration of longer-acting (*e.g*., pegylated) forms of IFN-λ .

Type I IFNs have been used to treat several viral diseases including chronic hepatitis C virus (HCV) and human papillomavirus (HPV) (Lazear et al., 2019). Although type I IFNs have garnered interest as a treatment strategy in COVID-19 (Palermo et al., 2021; Park and Iwasaki, 2020; Schreiber, 2020), their ability to exacerbate inflammation has tempered enthusiasm. One group tried to overcome this limitation by administering type I IFN by an intranasal route; in hamsters, they showed that nasally-delivered type I IFN could reduce viral burden, prevent virus transmission, and lower inflammation *in vivo* (Hoagland et al., 2021).

In our mouse models, administration of IFN-λ protected mice from infection, weight loss, lung inflammation, and lung disease, suggesting that the less pro-inflammatory nature of IFN-λ (Lazear et al., 2019) may have advantages as a therapeutic strategy. Our RNA sequencing data also showed IFN-λ treatment induced a tissue repair transcriptional signature in the lung, which contrasts with some studies showing that persistent type I or type III IFN signaling can disrupt lung epithelial barriers and prevent tissue repair (Broggi et al., 2020a;

Major et al., 2020). Nonetheless, administration of IFN-λ later in the course of SARS-CoV-2 infection, when most of the disease is caused by the host response and not by viral replication, could be detrimental and warrants further study.

By leveraging flow cytometry, qRT-PCR, and *Ifnl2-gfp* reporter mice, we found that IFN-λ was produced mainly from lung ECs after SARS-CoV-2 infection. This observation agrees with experiments by others after influenza A virus infection (Galani et al., 2017). We also showed IFN-λ acted primarily on radio-resistant cells in the lung to confer protection against SARS-CoV-2 infection, which is consistent with a recent finding (Broggi et al., 2020a). While others have suggested that IFN-λ signaling in neutrophils is required for optimal antifungal or antiviral defenses or limiting tissue damage (Broggi et al., 2017; Espinosa et al., 2017; Galani et al., 2017), our neutrophil depletion studies showed no effect on IFN-λ –mediated protection against SARS-CoV-2 infection or weight loss in mice. The basis for the difference is uncertain but could be due to the disparate models of pathogen infection or inflammation.

Although our experiments establish a role for IFN-λ in protecting against infection by SARS-CoV-2 strains including B.1.1.529, we acknowledge several limitations to our study:

(a) We used female mice in our IFN-λ treatment models, so studies in male animals are needed to exclude sex-based differences in therapeutic effects. Notwithstanding this, another group recently showed protective effects of IFN-λ against SARS-CoV-2-induced death in male K18-hACE2 mice (Sohn et al., 2021); (b) The relationship between induction of IFN-λ responses in mice and COVID-19 patients is unclear, especially given that many patients with severe disease have blunted IFN responses. While some of the diminished type I IFN response may be due to autoantibodies (Bastard et al., 2020; van der Wijst et al., 2021), the presence of such inhibitors against IFN-λ has not been described; (c) Although our neutrophil deletion and bone marrow chimera studies suggest that radio-resistant cells respond to IFN-λ to confer a protective antiviral effect, the precise cell type was not defined. Future studies with *Ifnlr1*^fl/fl^ conditional knockout mice are required to fully answer this question; and (d) Our studies are restricted to mice. IFN-λ treatment experiments in other animals (*e.g*., hamsters, ferrets, or nonhuman primates) and ultimately humans are needed for corroboration and determination of clinical utility.

In summary, we present evidence that nasal administration of IFN-λ confers pre- and post-exposure protection against infection by several SARS-CoV-2 strains including key variants of concern without causing extensive inflammation. In the lung, IFN-λ is induced in a MAVS and MyD88-dependent manner primarily in ECs that are likely infected, and acts upon radio-resistant cells to control infection. Additional treatment studies are warranted to evaluate further the potential of IFN-λ as a broadly-acting antiviral agent against SARS-CoV-2 and its emerging variants.

## Acknowledgements

This study was supported by grants and contracts from NIH: R01 AI157155, U01 AI151810, and 75N93019C00051 [all to M.S.D.], 75N93021C00014 [to Y.K.], F30 AI152327 [to E.S.W.], and T32GM139774-01 [to C.E.K.].

## Author Contributions

Z.C. and C.E.K. performed the infection experiments in mice.

Z.C. and E.S.W. titrated virus in tissues. Z.C. analyzed inflammation and pathology. J.Y. analyzed the RNAseq data. P.J.H. and Y.K. isolated and propagated the B.1.1.529 isolate.

M.S.D. obtained funding and supervised research. Z.C. and M.S.D. wrote the initial draft, with all other authors providing editorial comments.

## SUPPLEMENTAL FIGURE LEGENDS

**Figure S1. SARS-CoV-2 viral burden in infected K18-hACE2 mice, Related to Figure 2. (A**) Eight-week-old female K18-hACE2 mice were inoculated by intranasal route with 10^3^ FFU of WA1/2020 D614G. At -16 h before virus inoculation, mice were given 2 μg of murine IFN-λ2 or PBS by intraperitoneal injection. Viral RNA levels at 3 dpi (n = 6-7 per group, 2 experiments). (**B-G**) Eight-week-old female K18-hACE2 mice were inoculated by intranasal route with 10^3^ FFU of WA1/2020 D614G. At -16 h (**B-C**), D-3 (**D-E**) or +8 h (**F-G**), mice were given 2 μg of murine IFN-λ2 or PBS by intranasal route. Viral RNA (**B, D, and F**) and infectious virus (**C**, **E,** and **G**) levels at 3 dpi (**B-C**: n = 7 per group, 2 experiments; **D-E**: n = 8-9 per group, 2 experiments; **F-G**: n = 6-7 per group, 2 experiments). (**H-J**) Eight-week-old female K18-hACE2 mice were treated with 2 μ λ PBS by intranasal route at -16 h and +8 h relative to inoculation with 10^3^ FFU of WA1/2020 D614G and harvested at 7 dpi. (**H**) Weight change was monitored daily for 7 days. (**I**) Viral RNA levels at 7 dpi. (**J**) Infectious virus levels at 7 dpi (**H-J:** n = 9-10 per group, 2 experiments). Bars (**A-G and I-J**) indicate median values. Data were analyzed by Mann-Whitney test (**A-G and I-J**) or *t* tests of the area under the curve (**H**) (\**P* < 0.05, \*\**P* < 0.01, \*\*\**P* < 0.001, and \*\*\*\**P* < 0.0001).

**Figure S2. SARS-CoV-2 viral burden in the brains of K18-hACE2 and 129S2 mice, Related to Figures 2 and 3.** (**A-D**) Eight-week-old (**A-B**) or five-month-old (**C-D**) female K18-hACE2 mice were inoculated by intranasal route with 10^3^ FFU of WA1/2020 D614G (**A-B**) or B.1.1529 (**C-D**). At D-2 (**A**), D+1 and D+2 (**B and D**) or D-1 (**C**), mice were administered 2 μg of murine IFN-λ 2 or PBS by intranasal route. Viral RNA levels from brain at 3 dpi (**A**: n = 9 per group, 2 experiments; **B**: n = 8 per group, 2 experiments; **C**: n = 7-8 per group, 2 experiments; **D**: n = 6-7 per group, 2 experiments). (**E-H**) Six-week-old female 129S2 mice were inoculated by intranasal route with 10^5^ FFU of B.1.351. At D-1 (**E**), D-3 (**F**), D-5 (**G**) or -16 h and +8 h (**H**), mice were administered 2 μg of murine IFN-λ 2 or PBS by intranasal route. Viral RNA levels from brain at 4 dpi (**E**: n = 7 per group, 2 experiments; **F**: n = 6-8 per group, 2 experiments; **G**: n = 6-8 per group, 2 experiments; **H**: n = 8 per group, 2 experiments). Bars indicate median values. Data were analyzed by Mann-Whitney test (\*\**P* < 0.01 and \*\*\**P* < 0.001).

**Figure S3. Cytokine responses following IFN-λ treatment and SARS-CoV-2 infection, Related to Figure 2.** Eight-week-old female K18-hACE2 mice treated with 2 μg of murine IFN-λ 2 or PBS at -16 h by the intranasal route were challenged with 10^3^ FFU of WA1/2020 D614G. Cytokine levels in lung homogenates at 3 dpi (2 experiments, n = 7 per group except naïve n = 4). Data were analyzed by one-way ANOVA with Tukey’s multiple comparison test (\**P* < 0.05, \*\**P* < 0.01, \*\*\**P* < 0.001, and \*\*\*\**P* < 0.0001).

**Figure S4. Cytokine induction following IFN-λ treatment and SARS-CoV-2 infection, Related to Figure 3.** Six-week-old female 129S2 mice treated with two doses of 2 μg of murine IFN-λ2 or PBS at -16 h and +8 h by the intranasal route were challenged with 10^5^ FFU of B.1.351. Cytokine levels in lung homogenates at 4 dpi (n = 7 per group except naïve n = 4, 2 experiments). Data analyzed by one-way ANOVA with Tukey’s multiple comparison test (\**P* < 0.05, \*\**P* < 0.01, \*\*\**P* < 0.001 and \*\*\*\**P* < 0.0001).

**Figure S5. Heatmaps of RNA-seq data, Related to Figure 4.** Heatmaps of selected significantly upregulated or downregulated gene sets corresponding with IFN-λ2 treatment identified through GO analysis. Genes shown in each pathway are the union of the differentially expressed genes (DEGs) enriched in D+1 group or D+3 group versus control group (n = 4 per group). Columns represent sample groups and rows indicate genes.

**Figure S6. Flow cytometric gating strategy for lung cell populations, Related to Figure 5.** (**A-D**) For lung tissues, cells were gated on single, live, CD45^+^ and CD45^-^ cells. Alveolar macrophages (AM) were identified as CD45^+^ SiglecF^hi^ CD11c^hi^ cells, dendritic cells (DC) were identified as CD45^+^ SiglecF^-^ CD11c^+^ MHCII^+^ cells (**A**). B and T cells were identified as CD45^+^ CD19^+^ cells and CD45^+^ CD3^+^ cells, respectively (**B**). Neutrophils (Nϕ) and epithelial cells (EC) were identified as CD45^+^CD11b^+^Ly6G^+^ cells and CD45^-^ CD326^+^ cells, respectively (**C**). Monocytes (Mo) were identified as CD45^+^ CD11b^+^ Ly6C^hi^ cells (**D**).

**Figure S7. Flow cytometry analysis of peripheral blood from neutrophil-depleted or bone marrow chimeric mice, Related to Figure 6.** (**A**) Experimental scheme of neutrophil deletion in 129S2 mice. (**B**) (*Left*) Representative flow cytometry plots of peripheral blood at D+4 following intraperitoneal injection of a depleting anti-Ly6G mAb (1A8) or isotype control mAb. (*Right*) Frequency of mature neutrophils (CD11b^+^Ly6B^+^Ly6G^+^Ly6C^int^) in blood are shown after antibody depletion (n = 8 per group, 2 experiments). (**C**) Representative flow cytometry plots of peripheral blood at 10 weeks after irradiation and bone marrow cell transplantation of CD45.2 cells to CD45.1 recipient mice.

## STAR METHODS RESOURCE AVAILABILITY

### Lead contact

Further information and requests for resources and reagents should be directed to the Lead Contact, Michael S. Diamond.

### Materials availability

All requests for resources and reagents should be directed to the Lead Contact author. This includes mice, antibodies, viruses, and proteins. All reagents will be made available on request after completion of a Materials Transfer Agreement.

### Data and code availability

All data supporting the findings of this study are available within the paper and or upon request from the corresponding author. RNA sequencing datasets are available for analysis (GEO accession number GSE193990).

## EXPERIMENTAL MODEL AND SUBJECT DETAILS

### Cells and viruses

Vero-TMPRSS2 and Vero-TMPRSS2-ACE2 cells (Chen et al., 2021b) were cultured at 37[ in Dulbecco’s Modified Eagle medium (DMEM) supplemented with 10% fetal bovine serum (FBS), 10 mM HEPES pH 7.3, and 100 U/ml of penicillin–streptomycin. The SARS-CoV-2 WA1/2020 D614G virus was produced from an infectious clone and has been described previously (Chen et al., 2021b). The B.1.351 and B.1.1529 viruses were isolated from infected individuals and have been described previously (Chen et al., 2021a; VanBlargan et al., 2022). Infectious stocks were propagated in Vero-TMPRSS2 cells as described (Case et al., 2020). All work with infectious SARS-CoV-2 was performed in approved BSL3 and A-BSL3 facilities at Washington University School of Medicine using appropriate positive pressure air respirators and protective equipment.

### Mice

Animal studies were carried out in accordance with the recommendations in the Guide for the Care and Use of Laboratory Animals of the National Institutes of Health. The protocols were approved by the Institutional Animal Care and Use Committee at the Washington University School of Medicine. Virus inoculations were performed under anesthesia that was induced and maintained with ketamine hydrochloride and xylazine, and all efforts were made to minimize animal suffering. WT C57BL/6J (#000664) mice were obtained from The Jackson Laboratory or bred in a pathogen-free animal facility at Washington University. *Ifnlr1^-/-^* (Ank et al., 2008) and *Ifnl2-gfp* reporter mice (originally generated by Evangelos Andreakos, and kindly provided by Megan Baldridge, Washington University) were bred and housed in a pathogen-free animal facility at Washington University. Heterozygous K18-hACE C57BL/6J mice (strain: 2B6.Cg-Tg(K18-ACE2)2Prlmn/J) were obtained from The Jackson Laboratory. 129S2 mice were obtained from Charles River. Animals were housed in groups and fed standard chow diets.

## METHOD DETAILS

### Mouse infection, immune cell depletion and bone marrow chimeric mice studies

For neutrophil depletion, anti-Ly6G (BioXCell; clone 1A8) or an isotype control (BioXCell; clone 2A3) was administered to mice by intraperitoneal injection at D-1 (500 μg), D+1 (200 μg) and D+3 (200 μg) relative to B.1.351 inoculation. For bone marrow chimeric mice, six-week-old male and female WT (CD45.1) and *Ifnlr1^-/-^* (CD45.2) recipient mice were irradiated with 9 Gy (X-ray) total body irradiation (TBI). One day later, mice were injected with 5×10^6^ sex-matched bone marrow cells from donor WT (CD45.2) or *Ifnlr1^-/-^* (CD45.2) mice. Ten weeks later, peripheral blood cell from recipient chimeric mice were analyzed by flow cytometry as described below.

### Plaque assay

Vero-TMPRSS2-ACE2 cells were seeded at a density of 1.25 x 10^5^ cells per well in flat-bottom 24-well tissue culture plates. The following day, media was removed and replaced with 200 μL of 10-fold serial dilutions of sample, diluted in DMEM+2% FBS. One hour later, 1 mL of methylcellulose overlay was added. Plates were incubated for 72 h, then fixed with 4% paraformaldehyde (final concentration) in PBS for 1 hour. Plates were stained with 0.05% (w/v) crystal violet in 20% methanol and washed twice with distilled, deionized water. Plaques were counted, and titers were calculated according to a previously described method (Case et al., 2020).

### Measurement of viral RNA

Mice were euthanized and tissues were collected. Nasal washes were collected in 0.5 mL of PBS. Tissues were weighed and homogenized with zirconia beads in a MagNA Lyser instrument (Roche Life Science) in 1 mL of DMEM media supplemented with 2% FBS. Tissue homogenates were clarified by centrifugation at 10,000. Viral RNA from homogenized tissues or nasal washes was isolated using the MagMAX Viral RNA Isolation Kit (ThermoFisher) and measured by TaqMan one-step quantitative reverse-transcription PCR (RT-qPCR) on an ABI 7500 Fast Instrument. Copies of SARS-CoV-2 *N* gene RNA in samples were determined using a previously published assay (Case et al., 2020). Briefly, a TaqMan assay was designed to target a highly conserved region of the *N* gene (Forward primer: ATGCTGCAATCGTGCTACAA; Reverse primer: GACTGCCGCCTCTGCTC; Probe: /56-FAM/TCAAGGAAC/ZEN/AACATTGCCAA/3IABkFQ/). This region was included in an RNA standard to allow for copy number determination down to 10 copies per reaction. The reaction mixture contained final concentrations of primers and probe of 500 and 100 nM, respectively.

### Cytokine and chemokine protein measurements

Lung homogenates were incubated with Triton X-100 (1% final concentration) for 1 h at room temperature to inactivate SARS-CoV-2. Homogenates were analyzed for cytokines and chemokines by Eve Technologies Corporation (Calgary, AB, Canada) using their Mouse Cytokine Array/Chemokine Array 31-Plex (MD31) platform.

### Lung histology

Animals were euthanized before harvest and fixation of tissues. Lungs were inflated with 1.2 mL of 10% neutral buffered formalin using a 3-mL syringe and catheter inserted into the trachea. Tissues were embedded in paraffin, and sections were stained with hematoxylin and eosin. Images were captured using the Nanozoomer (Hamamatsu) at the Alafi Neuroimaging Core at Washington University.

### Flow cytometry analysis of peripheral blood

For analysis of immune cell depletion, peripheral blood cells were collected, and erythrocytes were lysed with ACK lysis buffer (GIBCO) and resuspended in RPMI supplemented with 10% FBS. Single cell suspensions were preincubated with Fc Block antibody (BD PharMingen) in PBS + 2% FBS for 10 min at room temperature before staining. Cells were incubated with antibodies against the following markers: BV421 anti-CD45, AF700 anti-Ly6C, FITC anti-Ly6B, PE-CY7 anti-Ly6G and APC anti-CD11b. All antibodies were used at a dilution of 1:200. Cells were stained for 20 min at 4 ℃, washed with PBS, fixed with 4% PFA for 15 min, washed with PBS and resuspended with FACS (PBS + 2% FBS + 2 mM EDTA) buffer.

### Lung digestion and cell sorting by flow cytometry

Lungs were collected and digested at 37 with 5 mg/mL of collagenase I (Worthington) and 1 mg/mL of DNase I (Roche) for 45 min in HBSS buffer. Digested lung tissues were minced, passed through a 40 μm strainer, and centrifuged at 500 g for 10 min. Red blood cells were lysed with ACK lysis buffer (GIBCO). Dead cells were removed by Dead Cell Removal Kit (STEMCELL) according manufacturer’s protocol. Single cell suspensions were incubated with APC-CY7 anti-CD45, APC anti-CD11b, BV421 anti-Ly6G, BV-421 anti-CD11c, PE anti-Siglec F (CD170), AF-700 anti-MHC II (I-A/I-E), BV421 anti-CD3, PE anti-CD19, APC anti-CD11b, PE anti-CD326 and PE anti-Ly6C antibodies as described above. AM (CD45^+^ SiglecF^hi^ CD11c^hi^), DCs (CD45^+^ SiglecF^-^ CD11c^+^ MHCII^+^), B cells (CD45^+^ CD19^+^), T cells (CD45^+^ CD3^+^), Nϕ (CD45^+^CD11b^+^Ly6G^+^), ECs (CD45^-^ CD326^+^) and Mo (CD45+ CD11b^+^ Ly6C^hi^) were sorted by flow cytometry (Sony SH800Sorter) under BSL3 conditions. RNA was extracted with RNeasy Micro Kit (QIAGEN) according to manufacturer’s protocol and then *Ifnl2*, *Ifnl3,* and *Ifnlr1* mRNA levels were measured by qRT-PCR as described above.

### Confocal microscopy

Lung tissues were collected as described above and fixed for 7 days. Tissues then were washed three time with PBS and placed into 30% sucrose in PBS overnight until sinking to the bottom of the tube. Tissues were placed into O.C.T. medium in cryomolds on dry ice, wrapped in aluminum foil, and stored in -80C. Sections were cut and embedded on superfrost glass slides. Slides were rinsed three times with PBS, blocked with 5% FBS, 1% BSA and 0.3% Triton X-100 in PBS, and incubated with rat anti-CD326 (1: 500), rabbit anti-nucleocapsid protein (1: 500), and chicken anti-GFP (1: 1000) primary antibodies at 4[ overnight. The next day, slides were stained with goat anti-chicken (1: 500), donkey anti-rabbit (1: 500) and donkey anti-rat (1: 500) secondary antibodies for 1 h at room temperature and with Hoechst dye (1:10000) for 5 min. Slides were washed with PBS once, mounted with AquaPoly, and stored in the dark at 4[ until imaged.

### RNA sequencing

RNA from lung tissues was extracted by RNeasy Pls Mini Kit (QIAGEN) according to manufacturer’s protocol. cDNA libraries were constructed starting with 10 ng of total RNA. cDNA was generated using the Seqplex kit (Sigma-Aldrich, St. Louis, MO) with amplification of 20 cycles. Library construction was performed using 100 ng of cDNA undergoing end repair, A-tailing, ligation of universal TruSeq adapters, and 8 cycles of amplification to incorporate unique dual index sequences. Libraries were sequenced on the NovaSeq 6000 (Illumina, San Diego, CA) targeting 40 million read pairs and extending 150 cycles with paired end reads. RNA-seq reads were aligned to the mouse Ensembl GRCh38.76 primary assembly with STAR program (version 2.5.1a) (Dobin et al., 2013). Gene counts were derived from the number of uniquely aligned unambiguous reads by Subread:featureCount (version 1.4.6-p5) (Liao et al., 2014). The ribosomal fraction, known junction saturation, and read distribution over known gene models were quantified with RSeQC (version 2.6.2) (Liao et al., 2014). All gene counts were preprocessed with the R package EdgeR (Robinson et al., 2010) to adjust samples for differences in library size using the trimmed mean of M values (TMM) normalization procedure. Viral and ribosomal genes and genes not expressed in at least five samples (the smallest group size) at a level greater than or equal to 1 count per million reads were excluded, resulting 19,280 unique genes in further analysis. The R package limma (Ritchie et al., 2015) with voomWithQualityWeights function (Liu et al., 2015) was utilized to calculate the weighted likelihoods for all samples, based on the observed mean-variance relationship of every gene and sample. Differentially expressed genes were defined as those with at least 2-fold difference between two individual groups at P < 0.05.

## QUANTIFICATION AND STATISTICAL ANALYSIS

Statistical significance was assigned when *P* values were < 0.05 using Prism version 8 (GraphPad). Tests, number of animals (n), median values, and statistical comparison groups are indicated in the Figure legends. Analysis of weight change was determined by *t* test or one-way ANOVA with Dunnett’s post-test of the area under the curve depending on the number of comparison groups. Viral burden was analyzed by Mann-Whitney test when comparing two groups, or one-way ANOVA or Kruskal-Wallis test with Dunnett’s post-test when comparing three or more groups. Cytokine data were analyzed by one-way ANOVA with Tukey’s multiple comparison test. qRT-PCR data were analyzed by one-way ANOVA with Dunnett’s post-test.

