## Supplementary figures and images for "Nasally-delivered interferon-λ protects mice against upper and lower respiratory tract infection of SARS-CoV-2 variants including Omicron"

### Figure S1

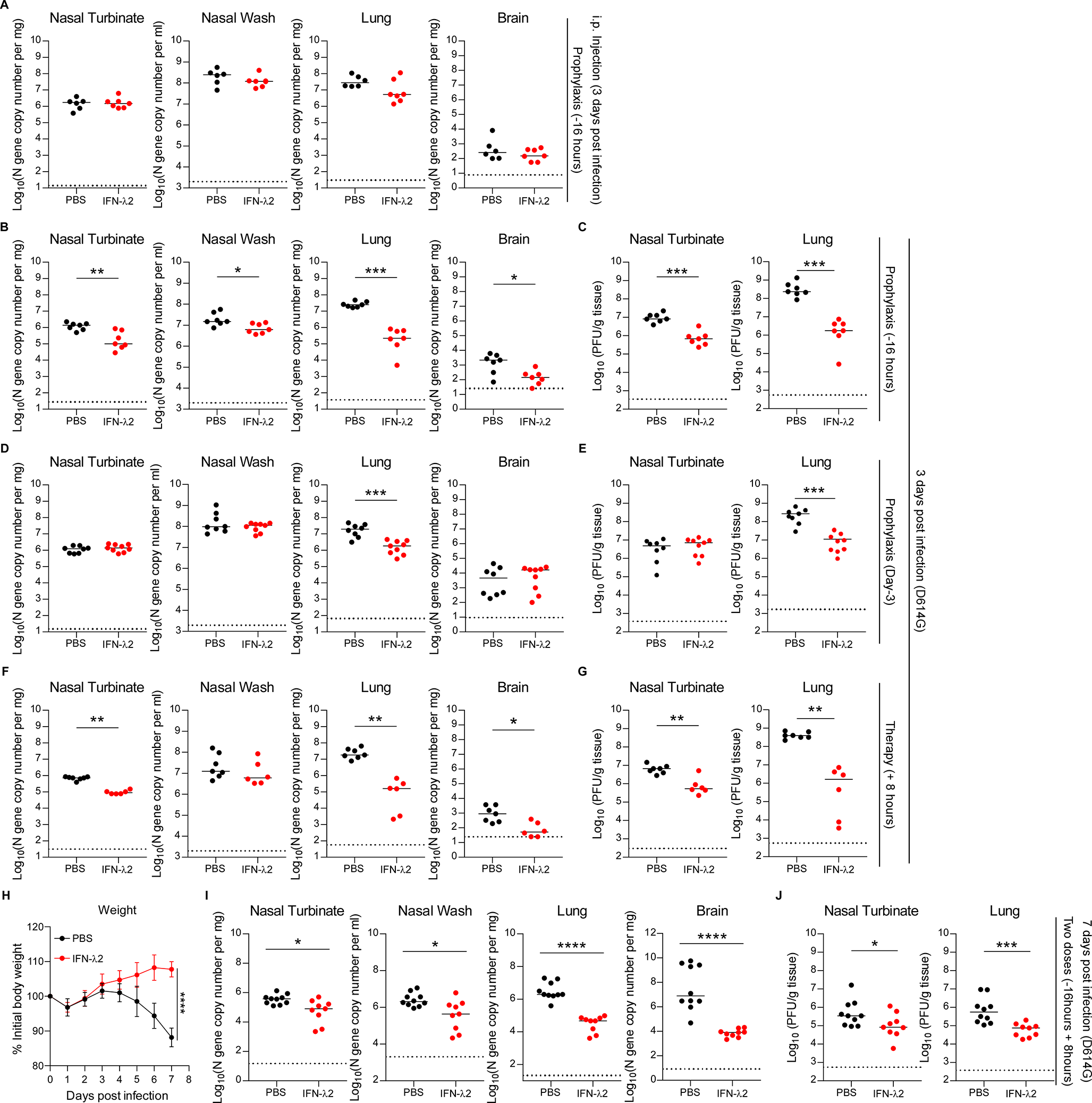

### Figure S2

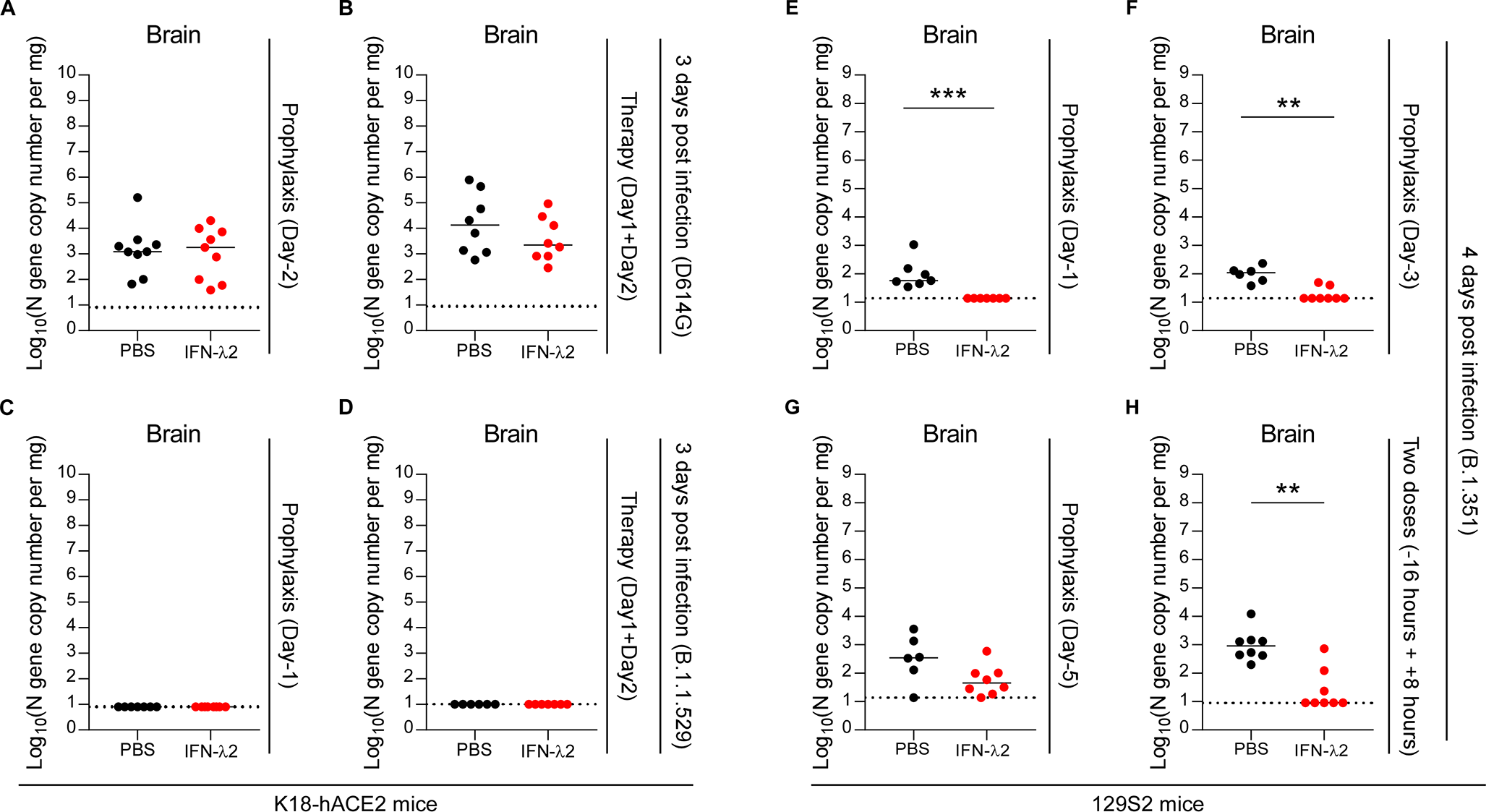

### Figure S3

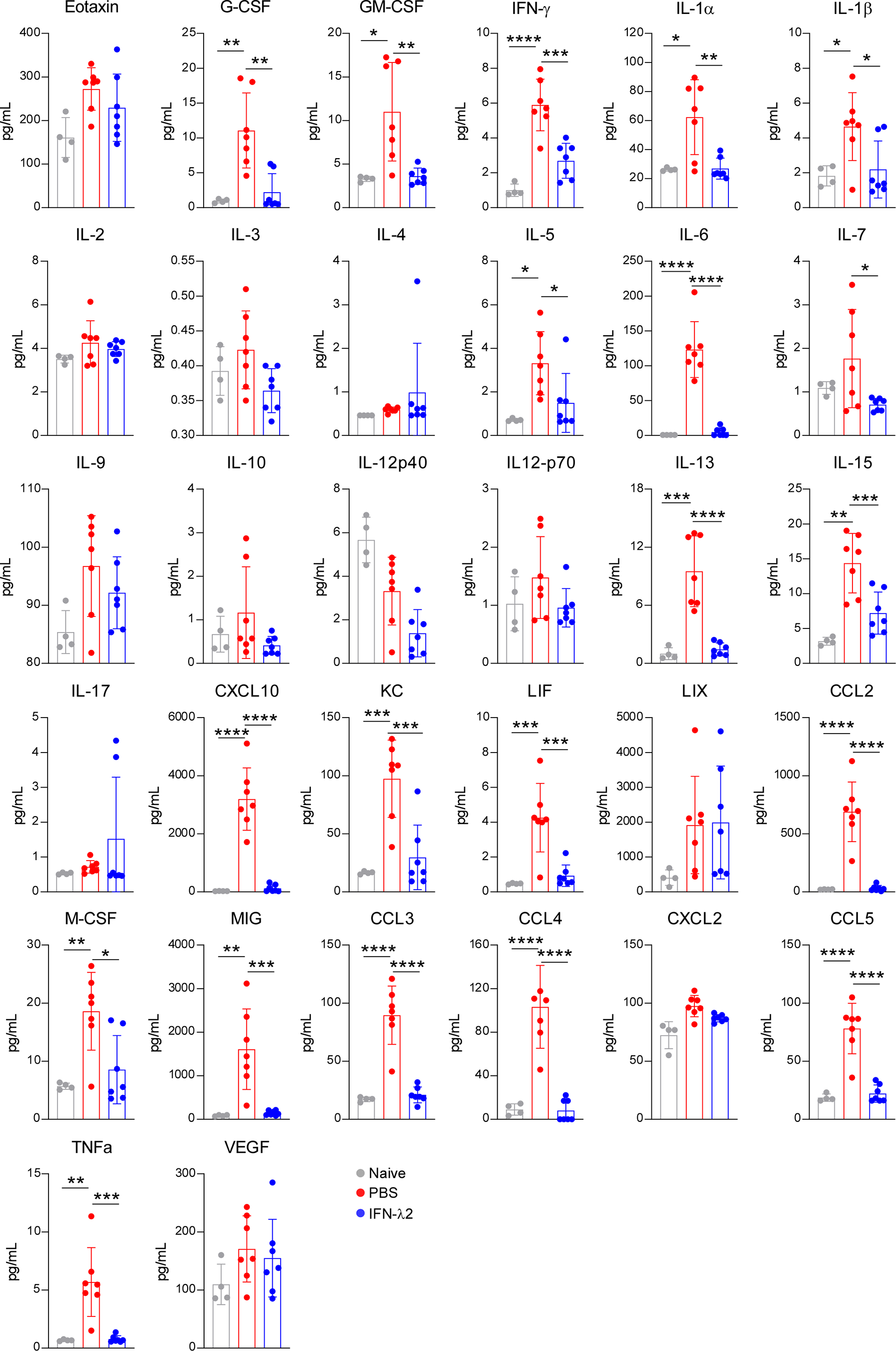

### Figure S4

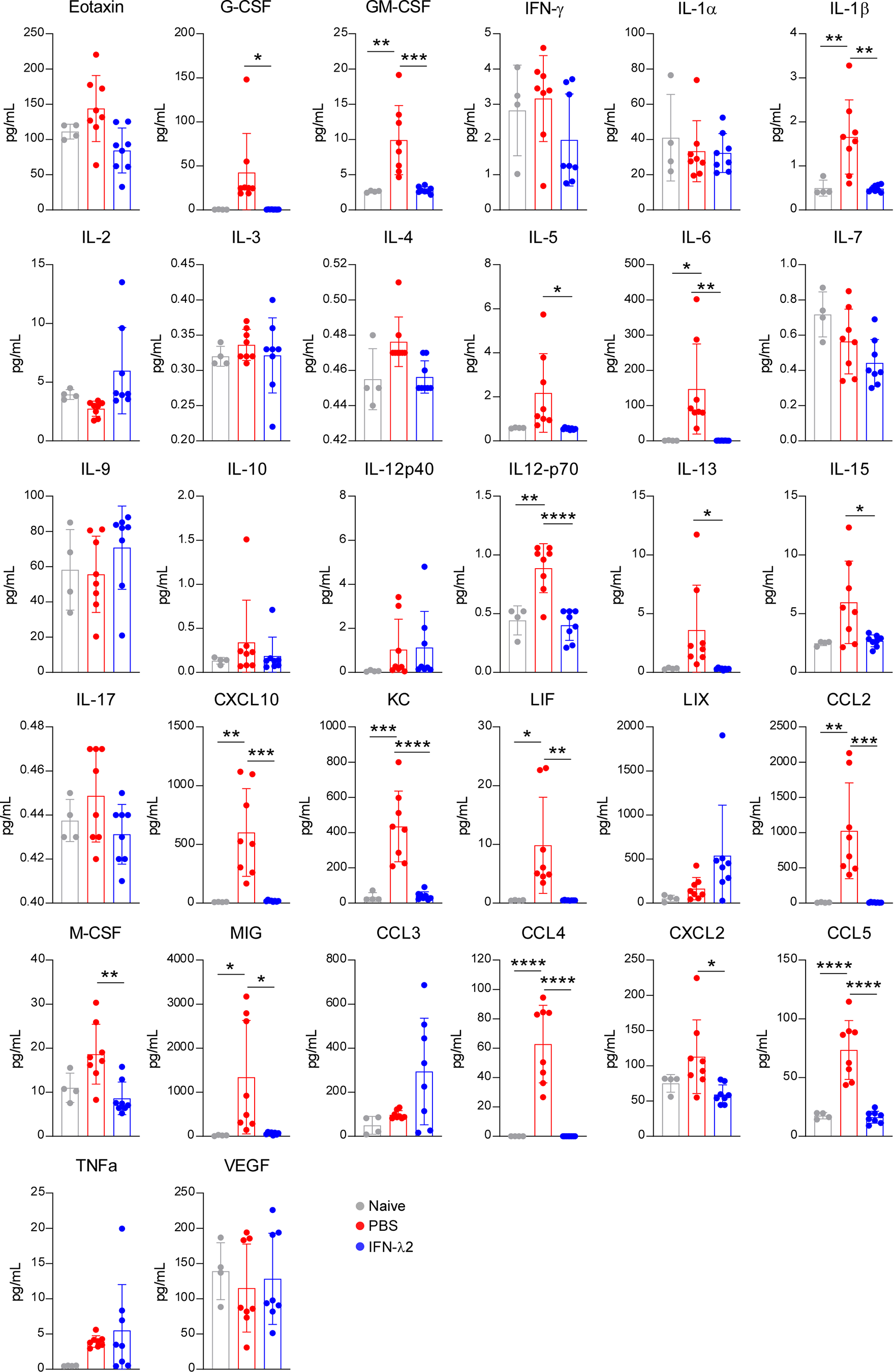

### Figure S5

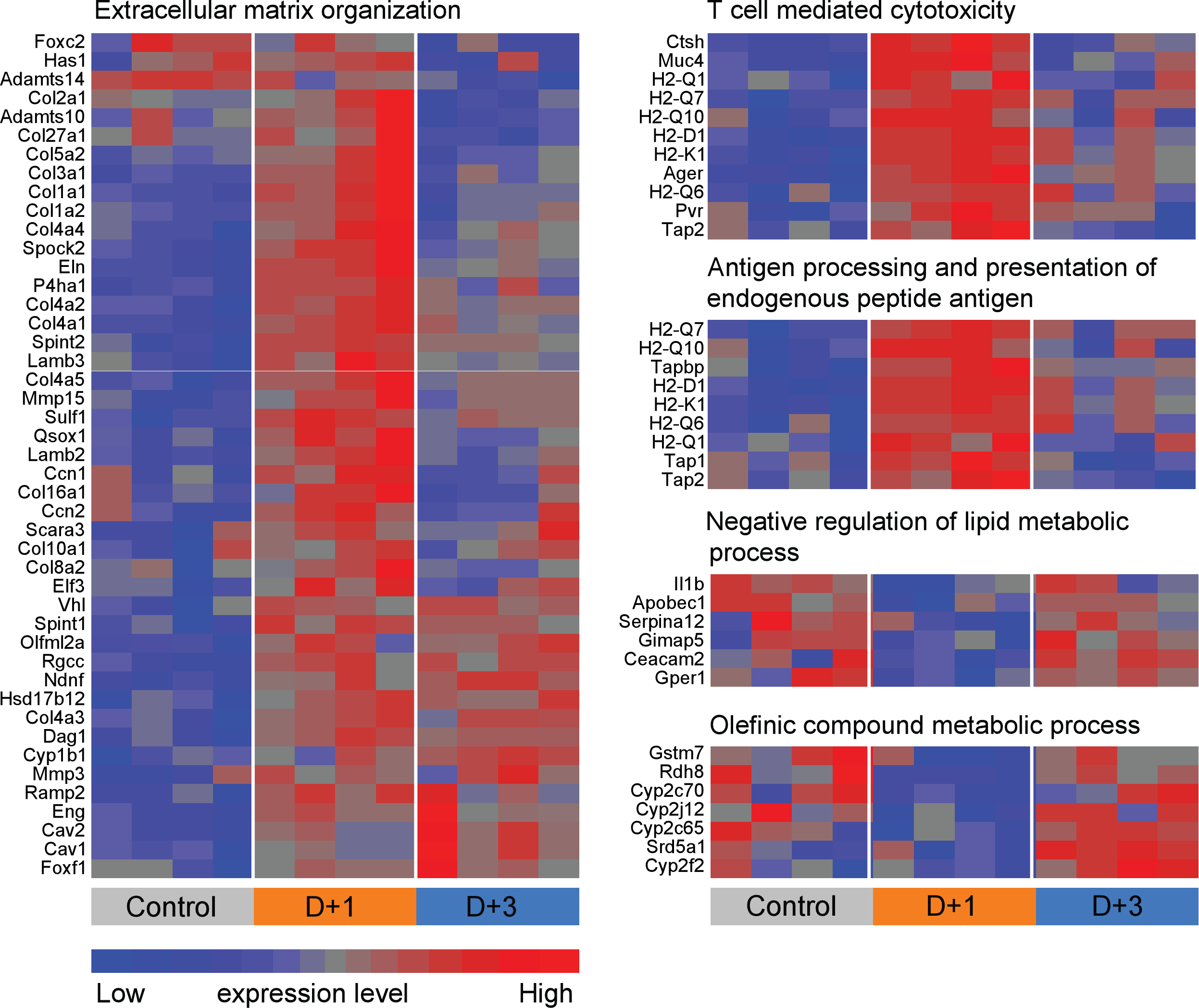

### Figure S6

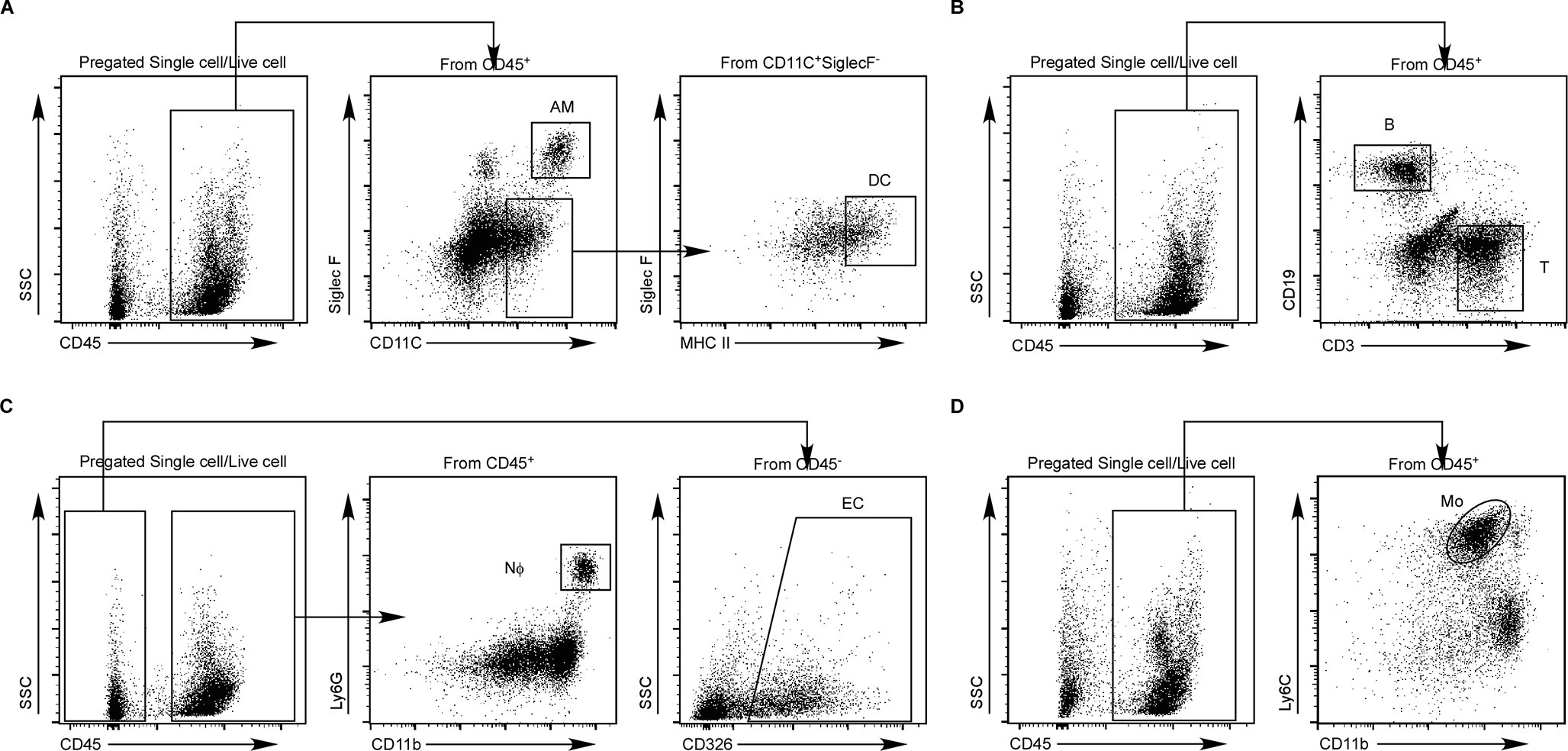

### Figure S7

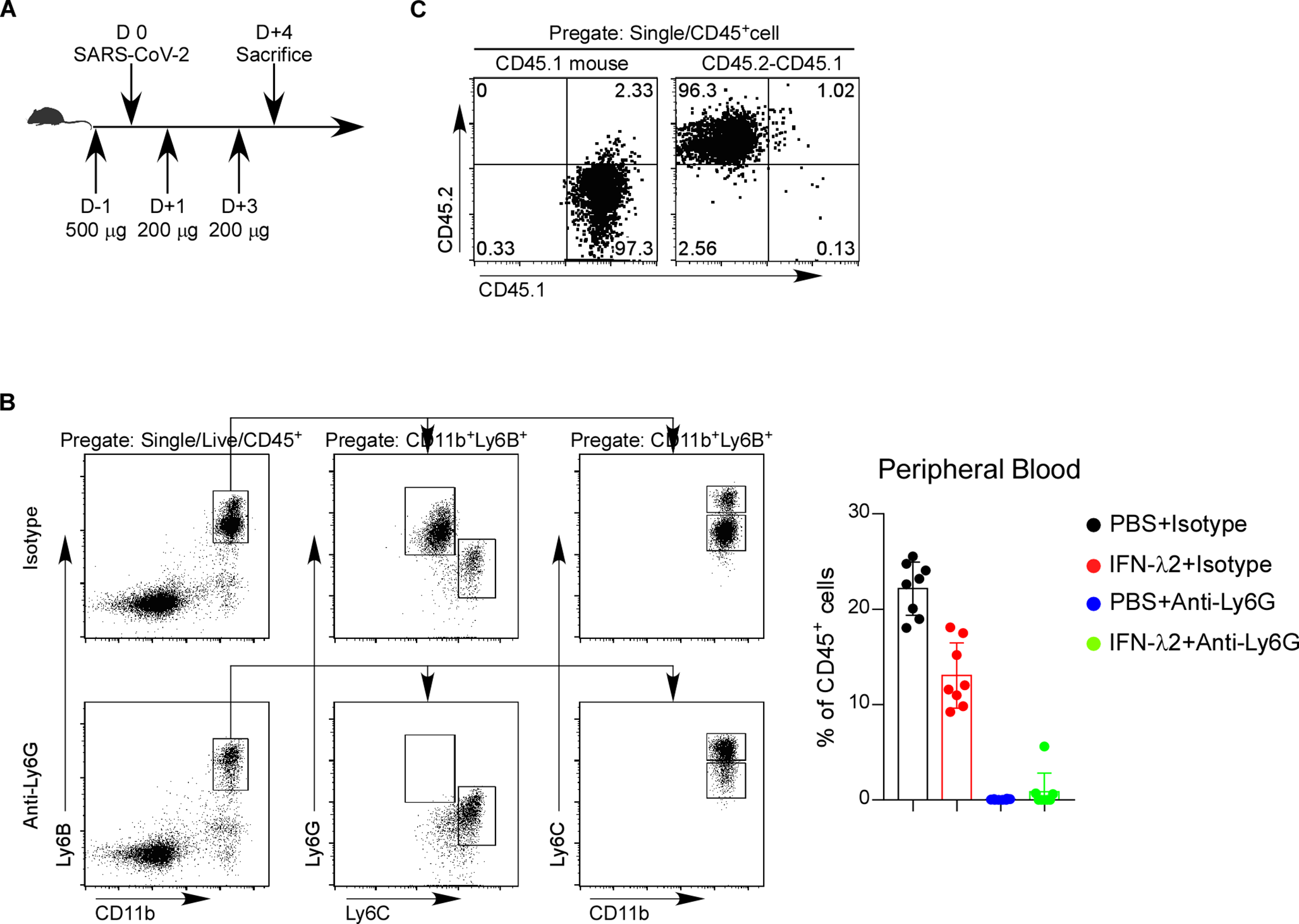
